# NFAT2 drives both Orai3 transcription and protein degradation by harnessing the differences in epigenetic landscape of MARCH8 E3 ligase

**DOI:** 10.1101/2024.12.11.628081

**Authors:** Sharon Raju, Akshay Sharma, Gyan Ranjan, Rajender K Motiani

## Abstract

The functional significance of any protein in physiological processes and pathological conditions is largely dependent on its expression profile. Therefore, nature has evolved several autonomous mechanisms to regulate protein expression such as transcription, translation, post-translational modifications and epigenetic changes. These processes are typically controlled by distinct molecular players with no overlapping roles. Here, we reveal that same transcription factor, NFAT2 regulates both transcription and lysosomal degradation of Orai3 oncochannel in a context dependent manner. We demonstrate that NFAT2 drives Orai3 transcription and thereby increases Orai3 levels in non-metastatic cancerous cells. While in invasive and metastatic cancerous cells, NFAT2 induces Orai3 lysosomal degradation by transcriptionally enhancing the levels of MARCH8 E3 ubiquitin ligase. Our biochemical and super-resolution microscopy data show that MARCH8 physically interacts with Orai3 eventually resulting in its degradation. Mechanistically, the dichotomy in regulation of Orai3 expression emerges from the differences in the epigenetic landscape of MARCH8. We uncover that the MARCH8 promoter is highly methylated in non-metastatic cancerous cells and hence NFAT2 does not induce MARCH8 mediated Orai3 degradation in these cells. Importantly, we demonstrate that MARCH8 restricts pancreatic cancer metastasis by targeting Orai3 degradation thereby highlighting pathophysiological importance of this signaling module. Taken together, we report a unique and clinically relevant scenario wherein nature has commissioned the same transcription factor to both enhance and curtail the expression of a target protein.

**Highlights:**

➢ NFAT2 transcriptionally upregulates Orai3 Ca^2+^ channel in non-metastatic cells
➢ NFAT2 induces Orai3 lysosomal degradation via MARCH8 E3 ubiquitin ligase in metastatic cells
➢ The dichotomy in NFAT2’s function is an outcome of differences in the methylation status of MARCH8 promoter
➢ MARCH8 inhibits pancreatic cancer metastasis by driving Orai3 degradation

**Graphical Abstract:** 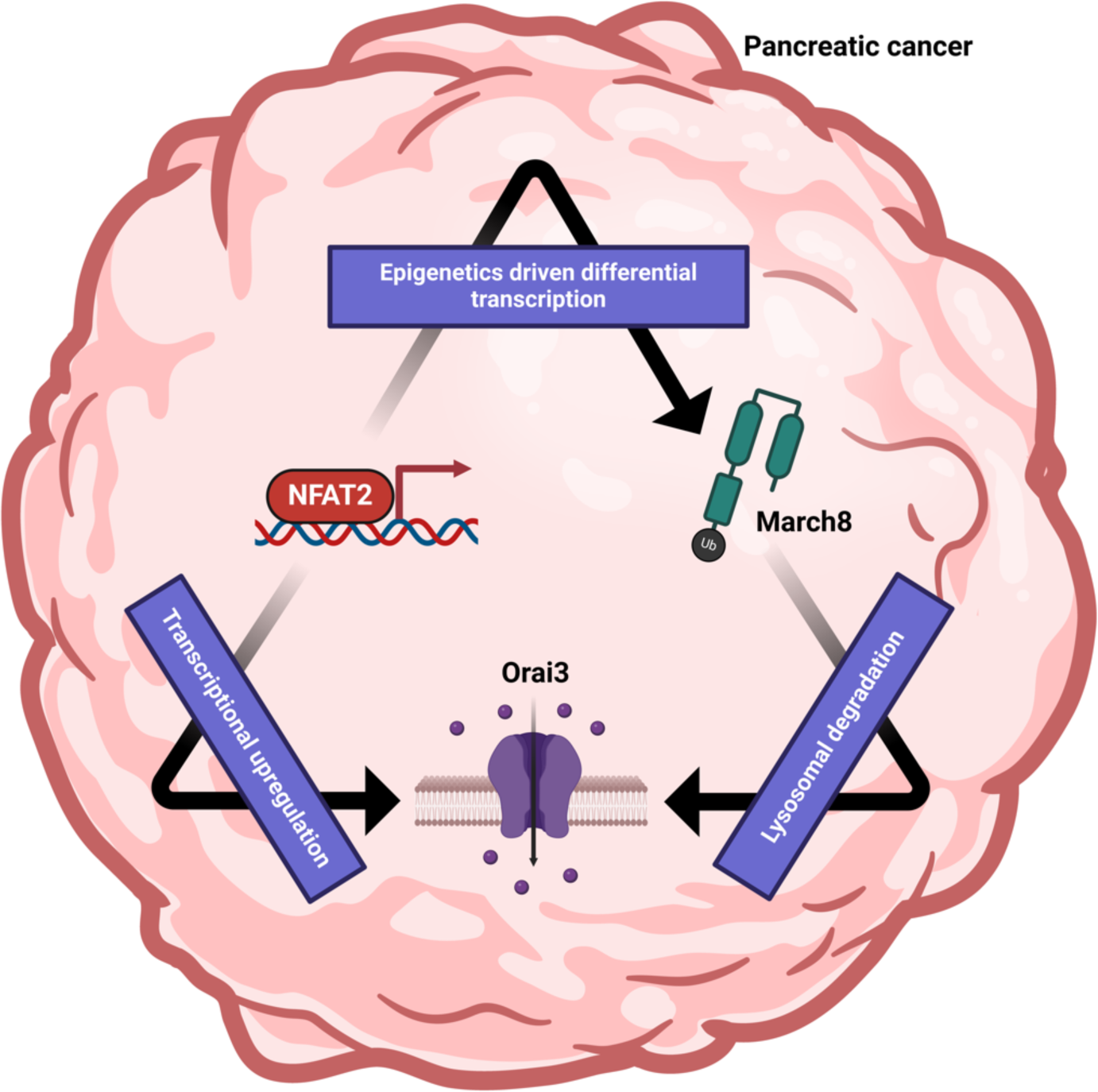

## Introduction

Proteins are involved in all aspects of cellular life from cell division to cell death. The regulation of proteins in the cell is a complex process involving various mechanisms that control when and how proteins are synthesized, modified, localized, and degraded (Ross et al., 2021). At the transcriptional level, gene expression regulates the amount of mRNA produced, while alternative splicing can lead to diverse protein variants. Post-transcriptionally, mRNA stability and translation are controlled by microRNAs and RNA-binding proteins. Once synthesized, proteins can undergo numerous post-translational modifications, such as phosphorylation, ubiquitination, and acetylation, which alter their activity, stability, and interactions. To prevent accumulation of damaged or excessive proteins, cells employ mechanisms like proteasome/lysosomal-mediated degradation and autophagy (Ross et al., 2021). Normally, each regulatory process for a single target protein is orchestrated by distinct molecular players with no intermingling effects on other mechanisms. The components involved in a specific protein regulatory process are typically limited to that process alone. These regulatory mechanisms work together to ensure precise spatio-temporal control over cellular functions and responses. But in pathophysiological conditions like cancer, proteome imbalance is a prominent feature due to alterations in these processes (Chen et al., 2023).

Pancreatic cancer (PC) is among the deadliest forms of cancer as it is highly metastatic in nature (Fitzmaurice et al., 2019). The 5-year survival rate of pancreatic cancer is around 13%, which is amongst the lowest. Ca^2+^ signaling plays a key role in regulating the oncogenesis and metastasis in cancer (Tanwar et al., 2020; Vashisht et al., 2015). However, functional relevance of Ca^2+^ dynamics in pancreatic cancer tumorigenesis is poorly understood. The store-operated calcium entry (SOCE) pathway is a ubiquitous process for Ca^2+^ influx into the cells. It is a process that starts with depletion of Ca^2+^ stores in the endoplasmic reticulum and results in Ca^2+^ influx across the plasma membrane. ER Ca^2+^ sensors STIM1/STIM2 sense the depletion of ER Ca^2+^ stores, oligomerize and move towards ER-plasma membrane junctions. STIM proteins physically interact with calcium-activated calcium release (CRAC) channels i.e Orai channels in the plasma membrane. This association activates Orai channels and Ca^2+^ influx across the plasma membrane (Kim et al., 2013; Lopez et al., 2020). Dysregulation in Orai channels in particular Orai3 expression and function is associated with various types of cancer (Chalmers & Monteith, 2018; Tanwar et al., 2020; Vashisht et al., 2015). However, the molecular mechanisms regulating Orai3 expression, stability and degradation remain largely unappreciated.

An earlier study from our group demonstrated that Orai3 forms a functional SOCE channel in pancreatic cancer cells and regulates key hallmarks of oncogenesis. Further, we demonstrated that Orai3 is transcriptionally upregulated in human pancreatic tumor samples as compared to normal patient samples (Arora et al., 2021). Pancreatic cancerous cells have higher Orai3 mRNA expression than normal cells, but what drives Orai3 transcriptional upregulation remains unappreciated. In addition to transcriptional regulation, protein degradation plays an essential role in regulating protein levels. Nevertheless, Orai3 degradation process remains completely unexplored. Gaining insight into the mechanisms that drive Orai3 protein degradation can aid in developing treatments for the diseases associated with aberrant Orai3 expression particularly pancreatic, breast, lung and prostate cancers.

Here, using unbiased and robust bioinformatic analysis, we found that Ca^2+^-sensitive NFAT2 transcription factor has putative binding sites on the Orai3 promoter. Interestingly, NFAT2 gain of function and loss of function studies revealed a dichotomy in the regulation of Orai3 by NFAT2 in non-metastatic versus invasive and metastatic cells. Our data demonstrates that NFAT2 positively regulates Orai3 transcription in non-metastatic pancreatic cancer cells. Hence, it generates a positive feedforward loop wherein a Ca^2+^-sensitive transcription factor drives transcription of a Ca^2+^ channel. Surprisingly, NFAT2 acts as a bimodal regulator in metastatic pancreatic cancer cells by inducing both Orai3 transcription and Orai3 protein degradation. Further, we show that downstream of NFAT2, MARCH8 E3 ubiquitin ligase ubiquitinates and degrades Orai3 via the lysosomal pathway. Mechanistically, NFAT2 transcriptionally regulates MARCH8 in pancreatic cancer cells depending on the epigenetic profile of the MARCH8 promoter. We reveal that the MARCH8 promoter is hypomethylated in invasive and metastatic pancreatic cancer cells compared to non-metastatic cells. Therefore, NFAT2 stimulates MARCH8 transcription in non-metastatic cells and thereby mediates Orai3 protein degradation. Finally, we establish that MARCH8 acts as a tumor suppressor in pancreatic cancer. Our *in vitro* migration assays and *in vivo* zebrafish xenograft experiments show that MARCH8 negatively regulates metastasis. To summarize, we reveal that same transcription factor controls both mRNA transcription and protein degradation of the identical target. Further, we have identified and characterized MARCH8 as an important regulator of Orai3 driven pancreatic cancer metastasis. Importantly, the dichotomy in Orai3 regulation highlights an intricate pathway in pancreatic cancer cells that can control disease outcomes.

## Results

### NFAT2 binds on the Orai3 promoter and regulates Orai3 expression

To delineate the transcriptional regulation of Orai3, we carried out extensive bioinformatics analysis of human Orai3 promoter through three independent tools: Eukaryotic promoter database (Perier, 2000), PSCAN (Zambelli et al., 2009), and Contra_V3 (Kreft et al., 2017). PSCAN was utilized to identify all potential transcription factor binding sites within the Orai3 promoter. PSCAN leverage the JASPAR Core 2020 transcription factor position weight matrix database. The analysis revealed that the NFAT family of transcription factors has potential binding sites on the Orai3 promoter (**Figure 1A**). To validate this observation, the Orai3 promoter was analyzed for NFATc-binding sites using the EPD-Search Motif Tool, applying a stringent p-value cut-off of p = 0.01. Moreover, we assessed the sequence of the Orai3 promoter for NFATc-binding sites using Contra_V3 with a similarity matrix value = 0.75 and core value = 0.90. All three tools, each using different algorithm, consistently identified three NFATc binding sites on the Orai3 promoter. The three NFATc binding sites were present at - 920bp, -990bp, and -1017bp before the transcription start site. Furthermore, the multi-species alignment of the Orai3 promoter showed that these binding sites were conserved across mammalian species (**Figure 1B, C**). Thus, our robust bioinformatics analysis suggests that NFAT transcription factors may transcriptionally regulate Orai3.

**Figure 1:**
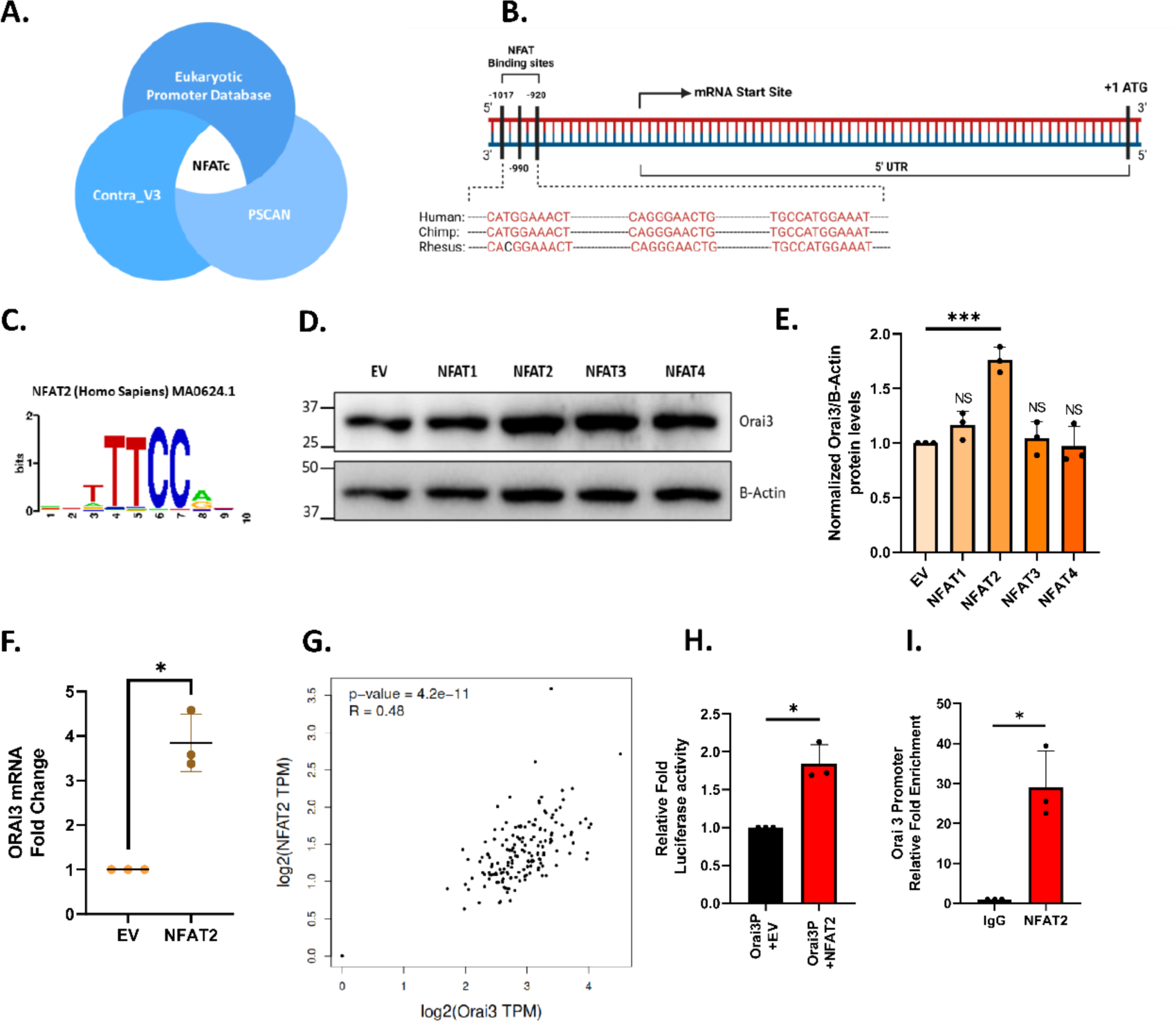
NFAT2 binds on Orai3 Promoter and regulates Orai3 expression. **A).** *In-silico* analysis via Eukaryotic Promoter Database, P-Scan and Contra_V3 establishing NFAT2 as a potential transcriptional regulator of Orai3. **B).** Identification of putative NFATc binding sites on the Human Orai3 promoter using the EPD-Search Motif Tool at p-val cut-off of 0.01 and bioinformatic characterization of conserved NFATc binding sites on the cross-species alignment of the Orai3 promoter using the ContraV3 transcription factor binding analysis tool. **C).** Position weight matrix for human NFAT2 consensus binding sequence. **D).** Representative western blot for Orai3 levels in transient overexpression of NFAT1, 2, 3 and 4 in HEK-293T cells. **E).** Densitometric analysis of Orai3 levels in NFAT1, 2, 3 and 4 overexpressed HEK-293T cells (N=3). **F).** qRT-PCR analysis showing transcriptional upregulation of Orai3 upon overexpression of NFAT2 in HEK-293T cells compared to control (N=3). **G).** NFAT2 and Orai3 mRNA expression analysis in GEPIA showing a positive correlation of NFAT2 and Orai3 in pancreatic tissues. **H).** Normalized Luciferase activity of Orai3 promoter in PANC-1 cells upon NFAT2 overexpression for 48 hrs. (N=3). **I).** ChIP-qPCR Analysis in PANC-1 cells for relative fold enrichment in NFAT2 immunoprecipitated DNA samples showing higher enrichment of Orai3 promoter region compared to IgG mock IP (N=3). Data presented are mean ± S.E.M. For statistical analysis, one sample *t*-test was performed for panels D, E, H and I using GraphPad Prism software. Here, NS means non-significant; * *p* <0.05 and *** *p* < 0.001.

The NFAT family of transcription factors has four Ca^2+^ sensitive members: NFAT1, 2, 3, and 4 (Müller & Rao, 2010). To identify which isoforms of NFAT regulate Orai3, we performed overexpression studies of the four isoforms of NFAT in HEK-293T cell line. We found that only NFAT2 significantly upregulates Orai3 expression (**Figure 1D, E**). Further, NFAT2 overexpression increased Orai3 mRNA levels compared to empty vector control (**Figure 1F**). To investigate the association of NFAT2 and Orai3 levels in pancreatic tissue, we carried out expression analysis of NFAT2 and Orai3 using the “GEPIA” (Gene Expression Profiling Interactive Analysis) database (Tang et al., 2017) and observed a positive correlation between NFAT2 and Orai3 mRNA expression in pancreatic tissue (**Figure 1G**).

To corroborate the bioinformatic analysis, we performed dual luciferase assays. Orai3 promoter was cloned into a luciferase reporter vector, and dual luciferase assays were performed with NFAT2 overexpression in PANC-1 pancreatic cancer cells. The results showed an increase in relative luciferase activity with NFAT2 overexpression compared to the empty vector control (**Figure 1H**) suggesting that NFAT2 positively regulates Orai3 expression. To determine if NFAT2 physically binds to the Orai3 promoter, we performed Chromatin Immunoprecipitation (ChIP) assays in PANC-1 cells either overexpressing eGFPC1-NFAT2 or an empty vector control (pEGFP-C1). The immunoprecipitated cross-linked sonicated DNA was amplified using human Orai3 promoter-specific primers. The ChIP-qPCR analysis showed that the Orai3 promoter with putative NFAT2 binding sites was highly enriched compared to mock IP samples (**Figure 1I**) thereby confirming physical binding of NFAT2 on Orai3 promoter. Collectively, the ChIP and dual luciferase assays provide substantial evidence for the binding of NFAT2 on the Orai3 promoter.

### NFAT2 overexpression dichotomously regulates Orai3 in non-metastatic v/s metastatic pancreatic cancer cells

To delineate the role of NFAT2 in regulating Orai3 expression and function, we chose three human pancreatic cancer cell lines: MiaPaCa-2 (non-metastatic), PANC-1 (invasive), and CFPAC-1 (metastatic). NFAT2 overexpression in these cell lines showed significant upregulation of Orai3 at mRNA level compared to empty vector control in non-metastatic cells (**Figure 2A**), invasive (**Figure 2F**) and metastatic cells (**Figure 2K**). However, at the protein level, we observed a dichotomy in Orai3 regulation by NFAT2. In non-metastatic cells, NFAT2 overexpression led to an increase in Orai3 at the protein levels (**Figure 2B, C**). Surprisingly, NFAT2 overexpression in invasive (**Figure 2G, H**) and metastatic cells (**Figure 2L, M**) led to a decrease in Orai3 protein levels compared to the control. This suggests that some sort of negative feedback loop is functional in invasive and metastatic cells. It appears that in these cells, NFAT2 apart from transcriptionally regulating Orai3, initiates a protein degradation cascade.

**Figure 2:**
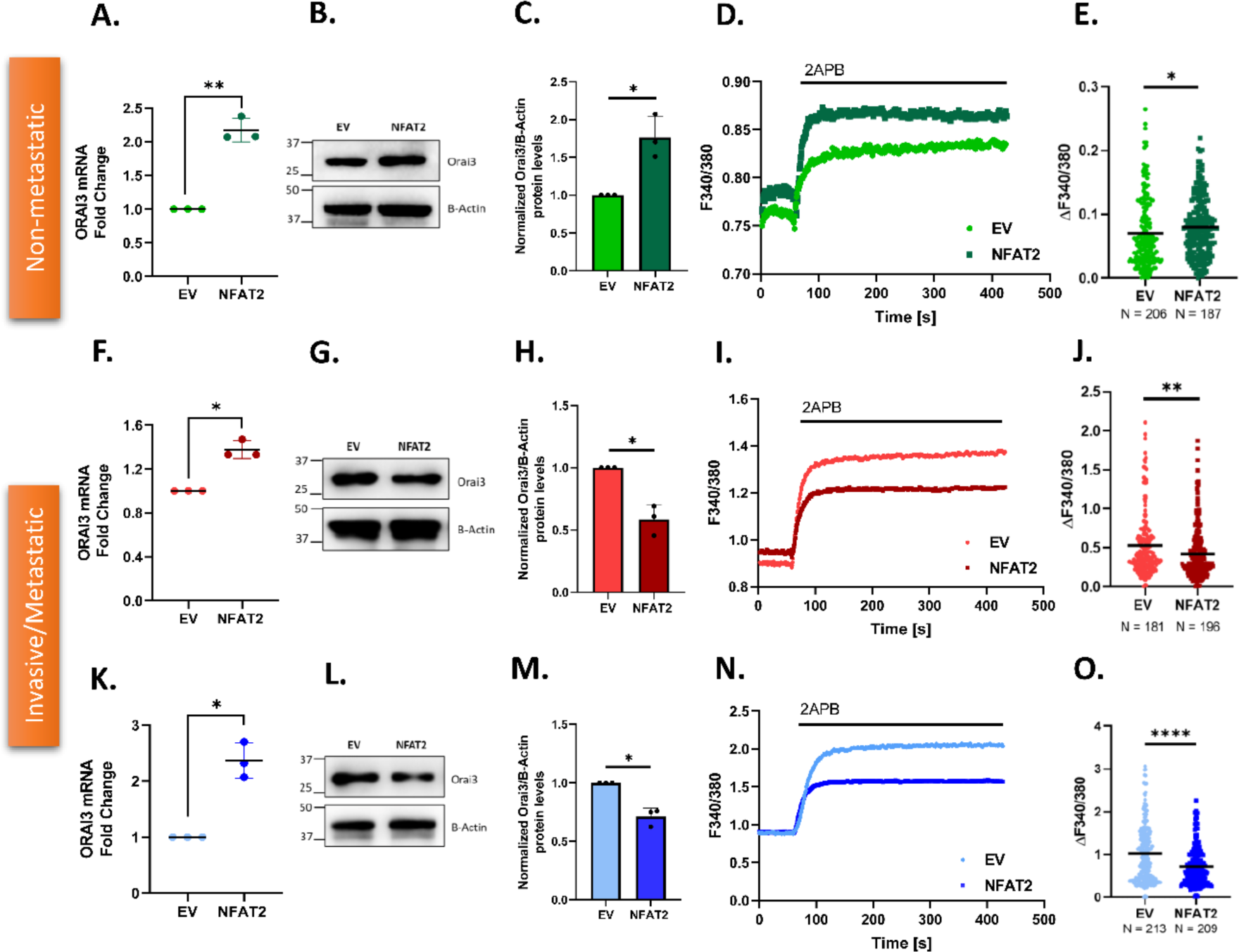
NFAT2 overexpression dichotomously regulates Orai3 in non-metastatic v/s metastatic PC cells. **A).** qRT-PCR analysis showing increase in Orai3 mRNA levels upon NFAT2 overexpression in MiaPaCa-2 compared to control (N=3). **B).** Representative western blots for increase in Orai3 protein levels due to NFAT2 overexpression in MiaPaCa-2 cells compared to control. **C).** Densitometric quantitation of Orai3 protein levels in NFAT2 overexpressed MiaPaCa-2 compared to control (N=3). **D).** Representative Ca^2+^ imaging trace of control vector pEGFP-C1 plasmid and NFAT2 overexpression plasmid in MiaPaCa-2. **E).** Orai3 potentiation of 2-APB in NFAT2 overexpressed and empty vector control MiaPaCa-2 where “n” denotes the number of ROIs. **F).** qRT-PCR analysis showing increase in Orai3 mRNA expression upon NFAT2 overexpression in PANC-1 compared to control (N=3). **G).** Representative western blots for decrease in Orai3 protein levels due to NFAT2 overexpression in PANC-1 compared to control. **H).** Western Blot densitometry of Orai3 protein levels in NFAT2 overexpressed PANC-1 cells compared to control (N=3). **I).** Representative Ca^2+^ imaging trace of control vector pEGFP-C1 plasmid and NFAT2 overexpression plasmid in PANC-1 cells. **J).** Potentiation of Orai3 by 2-APB in NFAT2 overexpressed and empty vector control PANC-1 where “n” denotes the number of ROIs. **K).** qRT-PCR analysis showing increase in Orai3 mRNA expression upon NFAT2 overexpression in CFPAC-1 compared to control (N=3). **L).** Representative western blots for decrease in Orai3 protein levels due to NFAT2 overexpression in CFPAC-1 compared to control. **M).** Western Blot densitometry of Orai3 protein in NFAT2 overexpressed CFPAC-1 cells compared to control (N=3). **N).** Representative Ca^2+^ imaging trace of control vector pEGFP-C1 plasmid and NFAT2 overexpression plasmid in CFPAC-1 cells. **O).** Potentiation of Orai3 by 2-APB in NFAT2 overexpressed and empty vector control CFPAC-1 where “n” denotes the number of ROIs. Data presented are mean ± S.E.M. For statistical analysis, one sample *t*-test was performed for panels A, C, F, H, K and M while unpaired student’s *t*-test was performed for panel D, I and N using GraphPad Prism software. Here, * *p* <0.05; ** *p* < 0.01 and **** *p* < 0.0001.

To further corroborate the dichotomous role of NFAT2 on Orai3, we examined Orai3 function upon NFAT2 overexpression in the three cell types. For this, we performed live-cell Ca^2+^ imaging with FURA-2AM dye, a ratio-metric Ca^2+^ indicator. We utilized 2-Aminoethoxydiphenyl borate (2APB), a pharmacological agent that can differentiate between functional Orai1 and Orai3 channels. It is widely recognized that 2APB, at concentrations ranging from 30 to 50 µM, inhibits Orai1 channels but activates Orai3 channels (Arora et al., 2021; Motiani et al., 2010, 2013; Vashisht et al., 2018). We utilized standard thapsigargin activated SOCE protocol (Tanwar et al., 2022, 2024). The Ca^2+^ imaging studies showed that in non-metastatic cells, NFAT2 overexpression led to an increase in ER Ca^2+^release (**Figure S1A, B**), augmentation in SOCE activity (**Figure S1A, C**) and higher 2APB potentiation (**Figure S1A, D**) compared to empty vector control. This data correlates with the protein data that NFAT2 overexpression upregulates Orai3 at protein level in non-metastatic cells. As reported earlier (Motiani et al., 2010), 2APB is an efficient tool to observe functional activity of Orai3, we did calcium imaging in HBSS-Ca buffer with 2APB addition alone. We observed that in non-metastatic cells, NFAT2 overexpression leads to higher 2APB-induced Orai3 potentiation compared to empty vector control (**Figure 2D, E**). This data correlates with both standard thapsigargin activated SOCE protocol calcium imaging and the Orai3 protein data that NFAT2 overexpression results in augmented Orai3 protein levels in these cells (**Figure 2B, C**). However, in invasive and metastatic cells, NFAT2 overexpression results in lower 2APB-induced Orai3 potentiation compared to empty vector control (**Figure 2I, J, N, O**). This functional analysis of Orai3 is in line with the protein data that NFAT2 overexpression downregulates Orai3 protein expression in invasive and metastatic cells. Ca^2+^ imaging with both standard thapsigargin activated SOCE protocol and 2APB potentiation replicated the same result. Therefore, for further imaging experiments, we continued with 2APB potentiation protocol.

### Competitive inhibition of NFAT validates dichotomous regulation of Orai3 in non-metastatic v/s metastatic PC cells

Overexpression of NFAT2 in PC cells indicated a bimodal regulation of Orai3 by NFAT2. To corroborate these findings, we inhibited NFAT activity with VIVIT. VIVIT is a small peptide that inhibits NFAT translocation to the nucleus by competing with NFATc for the calcineurin binding site (Aramburu et al., 1999). VIVIT peptide shows 25 times higher affinity for calcineurin binding and prevents calcineurin-facilitated NFAT dephosphorylation (Aramburu et al., 1999). Surprisingly, transfection of non-metastatic (Figure 3A), invasive (Figure 3F), and metastatic cells (Figure 3K) with VIVIT showed an increase in Orai3 at mRNA compared to empty vector control. One possible explanation for the increase in Orai3 mRNA by inhibiting NFAT in non-metastatic, invasive, and metastatic cells could be that other endogenous transcription factors are activated and they drive Orai3 transcription in this condition. To test this possibility, we performed luciferase assay in VIVIT transfected PANC-1 cells and observed an increase in relative luciferase activity in VIVIT transfected cells compared to empty vector control cells (**Figure S2A**). This indicates that inhibition of NFAT dephosphorylation activates other endogenous transcription factors to bind on Orai3 promoter and upregulate Orai3 transcription.

**Figure 3:**
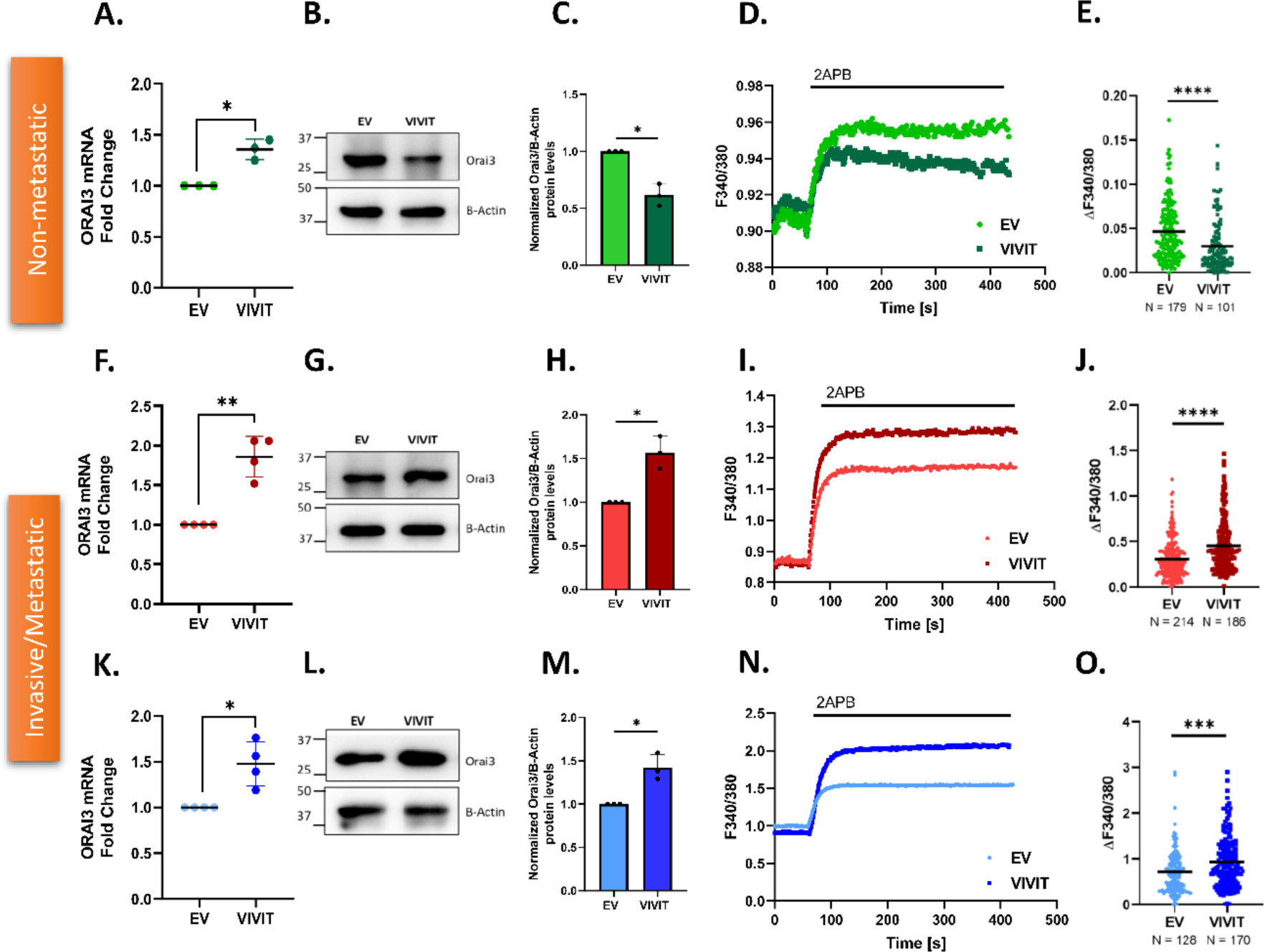
Competitive inhibition of NFAT validates dichotomous regulation of Orai3 in non-metastatic v/s metastatic PC cells. **A).** qRT-PCR analysis showing increase in Orai3 mRNA levels upon VIVIT transfection in MiaPaCa-2 compared to control (N=3). **B).** Representative western blots for decrease in Orai3 protein levels due to VIVIT transfection in MiaPaCa-2 cells compared to control. **C).** Densitometric quantitation of Orai3 protein levels VIVIT transfected MiaPaCa-2 compared to control (N=3). **D).** Representative Ca^2+^ imaging trace of control vector pEGFP-C1 plasmid and VIVIT transfection in MiaPaCa-2. **E).** Orai3 potentiation of 2-APB in VIVIT transfection and empty vector control MiaPaCa-2 where “n” denotes the number of ROIs. **F).** qRT-PCR analysis showing increase in Orai3 mRNA expression upon VIVIT transfection in PANC-1 compared to control (N=4). **G).** Representative western blots for increase in Orai3 protein levels due to VIVIT transfection in PANC-1 compared to control. **H).** Western Blot densitometry of Orai3 protein levels due to VIVIT transfection in PANC-1 cells compared to control (N=3). **I).** Representative Ca^2+^ imaging trace of control vector pEGFP-C1 plasmid and VIVIT transfection plasmid in PANC-1 cells. **J).** Potentiation of Orai3 by 2-APB in VIVIT transfected and empty vector control PANC-1 where “n” denotes the number of ROIs. **K).** qRT-PCR analysis showing increase in Orai3 mRNA expression upon VIVIT transfection in CFPAC-1 compared to control (N=4). **L).** Representative western blots for increase in Orai3 protein levels due to NFAT2 overexpression in CFPAC-1 compared to control. **M).** Western Blot densitometry of Orai3 protein levels in VIVIT transfected CFPAC-1 cells compared to control (N=3). **N).** Representative Ca^2+^ imaging trace of control vector pEGFP-C1 plasmid and VIVIT transfection plasmid in CFPAC-1 cells. **O).** Potentiation of Orai3 by 2-APB in VIVIT transfection and empty vector control CFPAC-1 cells where “n” denotes the number of ROIs. Data presented are mean ± S.E.M. For statistical analysis, one sample *t*-test was performed for panels A, C, F, H, K and M while unpaired student’s *t*-test was performed for panel D, I and N using GraphPad Prism software. Here, * *p* <0.05; ** *p* < 0.01; *** *p* < 0.001 and **** *p* < 0.0001.

Excitingly, NFAT2 inhibition by VIVIT resulted in a decrease in Orai3 protein expression in non-metastatic cells (Figure 3B**, C**) and an increase in Orai3 protein levels in invasive (Figure 3G**, H)** and metastatic cells (Figure 3L**, M**). This Orai3 protein data upon NFAT2 inhibition is in line with NFAT2 overexpression data. Collectively, the dichotomy in the regulation of Orai3 protein levels remains consistent in both NFAT2 gain of function and loss of function studies. Next, we performed standard thapsigargin activated SOCE protocol Ca^2+^ imaging in VIVIT-transfected non-metastatic and invasive cells. The Ca^2+^ imaging data revealed that inhibition of NFAT activity caused decrease in ER Ca^2+^ Release (**Figure S2B, C**), Ca^2+^ entry (**Figure S2B, D**) and 2APB potentiation (**Figure S2B, E**) in non-metastatic cells. This allied with the protein data in non-metastatic cells. Thus, NFAT inhibition decrease Orai3 activity in non-metastatic cells. Using 2APB mediated Orai3 potentiation protocol, we observed that inhibition of NFAT activity caused a decrease in 2APB-induced Orai3 potentiation in non-metastatic cells (Figure 3D**, E**). While in invasive (Figure 3I**, J**) and metastatic cells (Figure 3N**, O**), NFAT inhibition caused an increase in 2APB-induced Orai3 potentiation. Taken together, NFAT2 overexpression and inhibition data establishes the divergent control of NFAT2 over Orai3 protein expression and activity in non-metastatic v/s invasive and metastatic cells.

Since VIVIT inhibits transcriptional activity of all NFAT isoforms, we next validated the specificity of NFAT2 in the dichotomous regulation of Orai3. We performed siRNA mediated NFAT2 knockdown in pancreatic cancer cell lines (MiaPaCa-2, PANC-1 and CFPAC-1). We observed a robust decrease in the NFAT2 mRNA levels in all the cell lines **(Figure S2F, G, H)**. Next, we checked Orai3 mRNA and protein levels upon NFAT2 knockdown in these cells. Similar to VIVIT transfection, we observed an increase in Orai3 mRNA levels upon NFAT2 knockdown in MiaPaCa-2 **(Figure S2I)**, PANC-1 **(Figure S2N)** and CFPAC-1 **(Figure S2S)**. On the other hand, NFAT2 knockdown decreases Orai3 protein levels and functional activity in MiaPaCa-2 **(Figure S2J, K, L, M)** while Orai3 protein levels and functional activity increases upon NFAT2 silencing in PANC-1 **(Figure S2O, P, Q, R)** and CFPAC-1 **(Figure S2T, U, V, W)**. Therefore, both siRNA mediated NFAT2 knockdown and competitive inhibition of NFAT display the dichotomous regulation of Orai3 in non-metastatic v/s metastatic pancreatic cancer cells. Since NFAT2 siRNA and VIVIT gave same results, we proceeded with VIVIT for further NFAT2 loss of function studies.

### MARCH8 E3 Ubiquitin Ligase degrades Orai3

NFAT2 overexpression and inhibition studies in invasive and metastatic PC cell lines suggests involvement of Orai3 protein degradation in regulating Orai3 protein expression and activity. However, to best of our knowledge, the Orai3 protein degradation machinery is unknown. Therefore, we started investigating Orai3 protein degradation mechanism. First of all, to determine whether NFAT2-induced Orai3 protein degradation occurs via proteasomal or lysosomal pathway, we treated the NFAT2 overexpressed cells with either MG132 or bafilomycin A1. MG132 inhibits the proteasomal degradation pathway (Lee & Goldberg, 1998), while bafilomycin A1 blocks the lysosomal degradation cascade (Tapper & Sundler, 1995). We observed that treatment of bafilomycin A1 for 6hr in NFAT2 overexpressed invasive (Figure 4A**, B**) and metastatic cells (Figure 4C**, D**) inhibited Orai3 degradation by NFAT2 compared to the vehicle control. In contrast, treatment of MG132 for 6hr in NFAT2 overexpressed invasive (Figure 4A**, B**) and metastatic cells (Figure 4C**, D**) did not inhibit Orai3 degradation by NFAT2. This indicates that NFAT2 induces Orai3 degradation via lysosomal ubiquitination pathway in invasive and metastatic PC cells.

**Figure 4:**
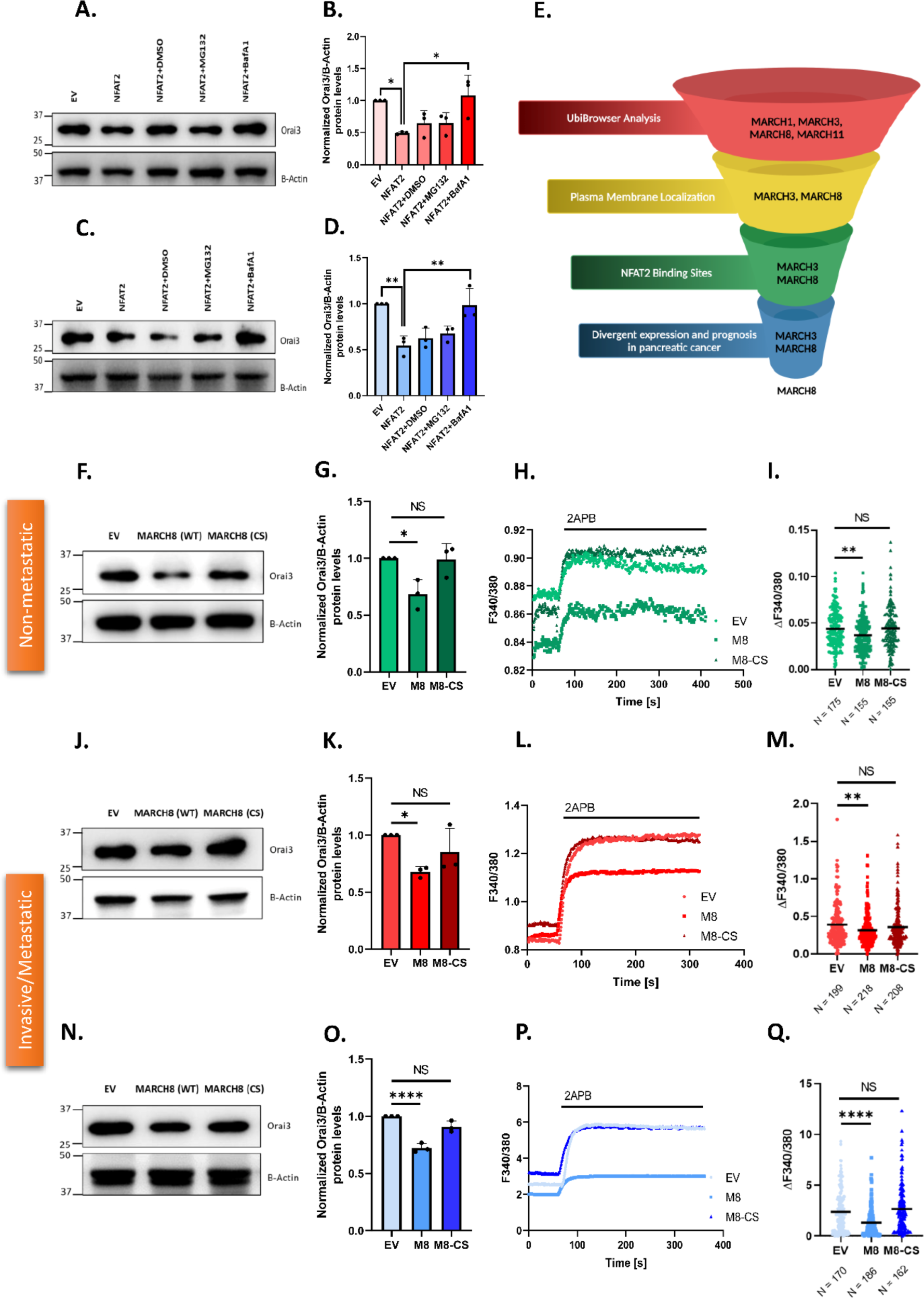
MARCH8 E3 Ubiquitin Ligase downregulates Orai3. **A).** Representative western blots for showing the effect of MG-132 (20 µM) and Bafilomycin-A1 (250nM) on NFAT2 induced protein degradation of Orai3 in PANC-1 cells. **B).** Densitometric analysis of Orai3 protein levels in NFAT2 overexpressed PANC-1 cells treated with MG-132 (20 µM) or Bafilomycin-A1 (250nM). **C).** Representative western blots for showing the effect of MG-132 (20 µM) and Bafilomycin-A1 (250nM) on NFAT2 induced protein degradation of Orai3 in CFPAC-1 cells. **D).** Densitometric analysis of Orai3 protein levels in NFAT2 overexpressed CFPAC-1 cells treated with MG-132 (20 µM) or Bafilomycin-A1 (250nM). **E).** Schematic of pipeline used for bioinformatic screening of potential E3 ubiquitin ligases targeting Orai3. **F).** Representative western blots showing overexpression of MARCH8 (WT) and not MARCH8 (CS) decreases Orai3 levels in MiaPaCa-2 cells compared to the control. **G).** Western Blot densitometry of Orai3 levels in MARCH8 (WT) and MARCH8 (CS) overexpressed MiaPaCa-2 cells (N=3). **H).** Representative Ca^2+^ imaging trace of control vector pEGFP-N1, MARCH8 (WT) and MARCH8 (CS) overexpression plasmids in MiaPaCa-2 cells. **I).** Potentiation of Orai3 by -APB in empty vector control, MARCH8 (WT) and MARCH8 (CS) overexpressed MiaPaCa-2 cells where “n” denotes the number of ROIs. **J).** Representative western blots showing overexpression of MARCH8 (WT) and not MARCH8 (CS) decreases Orai3 levels in PANC-1 cells compared to the control. **K).** Western Blot densitometry of Orai3 levels in MARCH8 (WT) and MARCH8 (CS) overexpressed PANC-1 cells (N=3). **L).** Representative Ca^2+^ imaging trace of control vector pEGFP-N1, MARCH8 (WT) and MARCH8 (CS) overexpression plasmids in PANC-1 cells. **M).** Potentiation of Orai3 by 2-APB in empty vector control, MARCH8 (WT) and MARCH8 (CS) overexpressed PANC-1 cells where “n” denotes the number of ROIs. **N).** Representative western blots showing overexpression of MARCH8 (WT) and not MARCH8 (CS) decreases Orai3 levels in CFPAC-1 cells compared to the control. **O).** Western Blot densitometry of Orai3 levels in MARCH8 (WT) and MARCH8 (CS) overexpressed CFPAC-1 cells (N=3). **P).** Representative Ca^2+^ imaging trace of control vector pEGFP-N1, MARCH8 (WT) and MARCH8 (CS) overexpression plasmids in CFPAC-1 cells. **Q).** Potentiation of Orai3 by 2-APB in empty vector control, MARCH8 (WT) and MARCH8 (CS) overexpressed CFPAC-1 cells where “n” denotes the number of ROIs. Data presented are mean ± S.E.M. For statistical analysis, ordinary one-way ANOVA was performed followed by Tukey’s multiple comparison test for panels B, D, G, I, K, M, O and Q using GraphPad Prism software. Here, NS means non-significant; * *p* <0.05; ** *p* < 0.01; and **** *p* < 0.0001.

As ubiquitination of proteins is mainly carried out by E3 ubiquitin ligases, we next started identifying the putative E3 ubiquitin ligases that target Orai3. We utilized well established bioinformatics tool (UbiBrowser) (X. Wang et al., 2022) to predict potential Orai3 E3 ubiquitin ligases. We only got 4 putative hits through this analysis. Interestingly, all the 4 predicted E3 ubiquitin ligases belong to the MARCH (Membrane-associated RING-CH-type) family of membrane bound E3 ubiquitin ligases i.e. MARCH1, MARCH3, MARCH8 and MARCH11.

Since Orai3 is a plasma membrane Ca^2+^ channel, its ubiquitination by membrane bound E3 ubiquitin ligases appears logical. MARCH family are a subset of RING domain containing E3 ubiquitin ligases, which have specific transmembrane domains allowing them to be tethered to the plasma membrane or membranes of other organelles. We next evaluated the subcellular localization of the 4 predicted E3 ubiquitin ligases. Intriguingly, only MARCH3 and 8 have been reported to be localized at the plasma membrane (Lin et al., 2019) while MARCH1 and MARCH11 are localized to endo-lysosomes. In light of this analysis, we then focused our attention on MARCH3 and MARCH8. Since we observed Orai3 lysosomal degradation downstream of NFAT2, we examined the presence of potential NFAT2 binding sites on the MARCH3 and MARCH8 promoters. Interestingly, we found that the promoters of both MARCH3 and MARCH8 have NFAT2 binding sites **(Figure S3A)**. As Orai3 is overexpressed in pancreatic cancer tissue and is associated with poor prognosis (Arora et al., 2021) we next analyzed the expression of MARCH3 and MARCH8 in pancreatic cancer samples **(Figure S3B, C).** Further, we examined association of MARCH3 and MARCH8 with pancreatic cancer patient survival using OncoLnc database (Anaya, 2016). Interestingly, we found that MARCH8 but not MARCH3 is associated with favorable prognosis in pancreatic cancer tissue wherein higher MARCH8 expression correlated with better patient survival **(Figure S3D, E)**. Additionally, MARCH8 is reported to have anti-oncogenic effects in breast and lung cancer (W. Chen et al., 2021; Qian et al., 2021). Moreover, MARCH8 promotes lysosomal degradation of target proteins (W. Chen et al., 2021). Taken together, this robust and extensive bioinformatics analysis suggested that MARCH8 could be the E3 ubiquitin ligase that bridges NFAT2 and Orai3 protein degradation (Figure 4E). Therefore, we decided to examine the role of MARCH8 in Orai3 degradation downstream of NFAT2.

To study the effect of MARCH8 on Orai3 protein levels, we overexpressed MARCH8 (WT) and inactive MARCH8 (CS) mutant in MiaPaCa-2 (non-metastatic), PANC-1 (invasive) and CFPAC-1 (metastatic) cells. The MARCH8 (CS) mutant is inactive as all the cysteine residues (Cys80, 83, 97, 99, 110, 123, and 126) are substituted with serine residues (Fujita et al., 2013). The overexpression of MARCH8 leads to decrease in Orai3 protein levels in non-metastatic (Figure 4F**, G**), invasive (Figure 4J**, K**), and metastatic cells (Figure 4N**, O**) while overexpression of MARCH8 (CS) mutant did not alter Orai3 protein levels in these cells. Next, Ca^2+^imaging upon MARCH8 (WT) and MARCH8 (CS) overexpression in MiaPaCa-2, PANC-1, and CFPAC-1 cells showed decrease in 2APB induced Orai3 potentiation upon MARCH8 (WT) overexpression but no significant change in 2APB induced Orai3 potentiation upon MARCH8 (CS) overexpression in non-metastatic (Figure 4H**, I**), invasive (Figure 4L**, M**), and metastatic cells (Figure 4P**, Q**). The Ca^2+^ imaging data further substantiated the Orai3 protein data. Taken together, this demonstrates that overexpression of functional MARCH8 leads to decrease in Orai3 expression and subsequent activity.

To further validate the role of MARCH8 in regulating Orai3 expression, we performed siRNA-mediated knockdown of MARCH8. We validated the knockdown of MARCH8 upon siMARCH8 transfections at mRNA and protein levels in MiaPaCa-2 (**Figure S4A, B**) PANC-1 (**Figure S4C, D**), CFPAC-1 (**Figure S4D, E**). We next looked into Orai3 protein levels and observed that MARCH8 silencing leads to increase in Orai3 protein levels in MiaPaCa-2 (Figure 5A**, B)**, PANC-1 (Figure 5E**, F)**, and CFPAC-1 (Figure 5I**, J)** cell lines compared to empty vector control. Subsequently, Ca^2+^ imaging in siRNA-mediated MARCH8 knockdown cells showed an increase in 2APB-induced Orai3 potentiation in MiaPaCa-2 (Figure 5C**, D)**, PANC-1 (Figure 5G**, H)**, and CFPAC-1 (Figure 5K**, L)** cells compared to siNT control cells. Hence, the Orai3 protein levels and activity is exactly opposite to the MARCH8 expression. Collectively, MARCH8 gain of function, overexpression of inactive MARCH8 and MARCH8 loss of function studies reveal that MARCH8 E3 ubiquitin ligase is a critical regulator of Orai3 protein levels.

**Figure 5:**
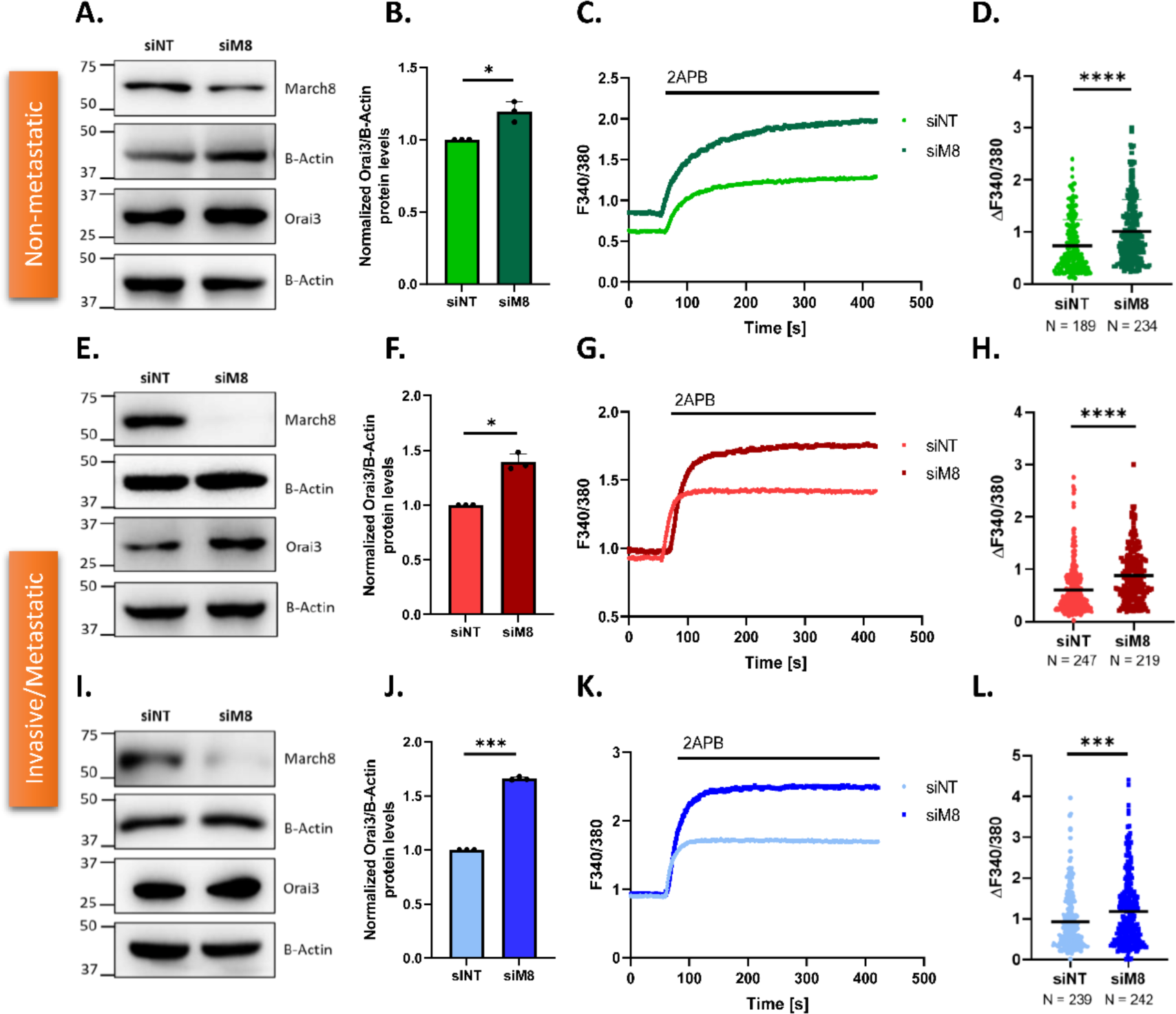
MARCH8 knockdown upregulates Orai3. **A).** Representative western blots showing increase in Orai3 protein levels upon siRNA mediated MARCH8 knockdown in MiaPaCa-2 cells compared to the control. **B).** Western Blot densitometry of Orai3 levels in siNT and siM8 transfected MiaPaCa-2 cells (N=3). **C).** Representative Ca^2+^ imaging trace of siNT and siM8 transfected MiaPaCa-2 cells. **D).** Potentiation of Orai3 by 2-APB in siNT and siM8 transfected MiaPaCa-2 cells where “n” denotes the number of ROIs. **E).** Representative western blots showing increase in Orai3 protein levels upon siRNA mediated MARCH8 knockdown in PANC-1 cells compared to the control. **F).** Western Blot densitometry of Orai3 levels in siNT and siM8 transfected PANC-1 cells (N=3). G). Representative Ca^2+^ imaging trace of siNT and siM8 transfected PANC-1 cells. **H).** Potentiation of Orai3 by 2-APB in siNT and siM8 transfected PANC-1 cells where “n” denotes the number of ROIs. **I).** Representative western blots showing increase in Orai3 protein levels upon siRNA mediated MARCH8 knockdown in CFPAC-1 cells compared to the control. **J).** Western Blot densitometry of Orai3 levels in siNT and siM8 transfected CFPAC-1 cells (N=3). **K).** Representative Ca^2+^ imaging trace of siNT and siM8 transfected CFPAC-1 cells. **L).** Potentiation of Orai3 by 2-APB in siNT and siM8 transfected CFPAC-1 cells where “n” denotes the number of ROIs. Data presented are mean ± S.E.M. For statistical analysis, one sample t-test was performed for panels B, D, F, H, J and L using GraphPad Prism software. Here, * *p* <0.05; *** *p* < 0.001; and **** *p* < 0.0001.

### MARCH8 physically interacts with Orai3 and induces its ubiquitination

MARCH8 E3 ubiquitin ligase ubiquitinates its target protein at its lysine residues, which leads to target degradation. Hence, we performed ubiquitination assays to check the status of Orai3 ubiquitination by immunoprecipitating Orai3 using an antibody against Orai3 followed by immunoblotting with ubiquitin antibody. As shown in Figure 6A, the immunoprecipitated Orai3 shows ubiquitination. Next, we investigated the role of MARCH8 in Orai3 ubiquitination. We overexpressed MARCH8 in bafilomycin-A1 treated PANC-1 cells and performed Orai3 immunoprecipitation. The western blot analysis shows that MARCH8 overexpression increases Orai3 ubiquitination compared to the empty vector control (Figure 6B). This data suggests that MARCH8 E3 ubiquitin ligase ubiquitinates Orai3. To study the interaction of MARCH8 and Orai3, we performed immunofluorescence assays and found that Orai3 and MARCH8 colocalize in the plasma membrane (Figure 6C**, D**). Further, we carried out co-immunoprecipitation assays and observed that MARCH8 and Orai3 interact with each other (Figure 6E). Together, biochemical and microscopy data demonstrate that MARCH8 physically interacts with Orai3 at plasma membrane and induces Orai3 ubiquitination.

**Figure 6:**
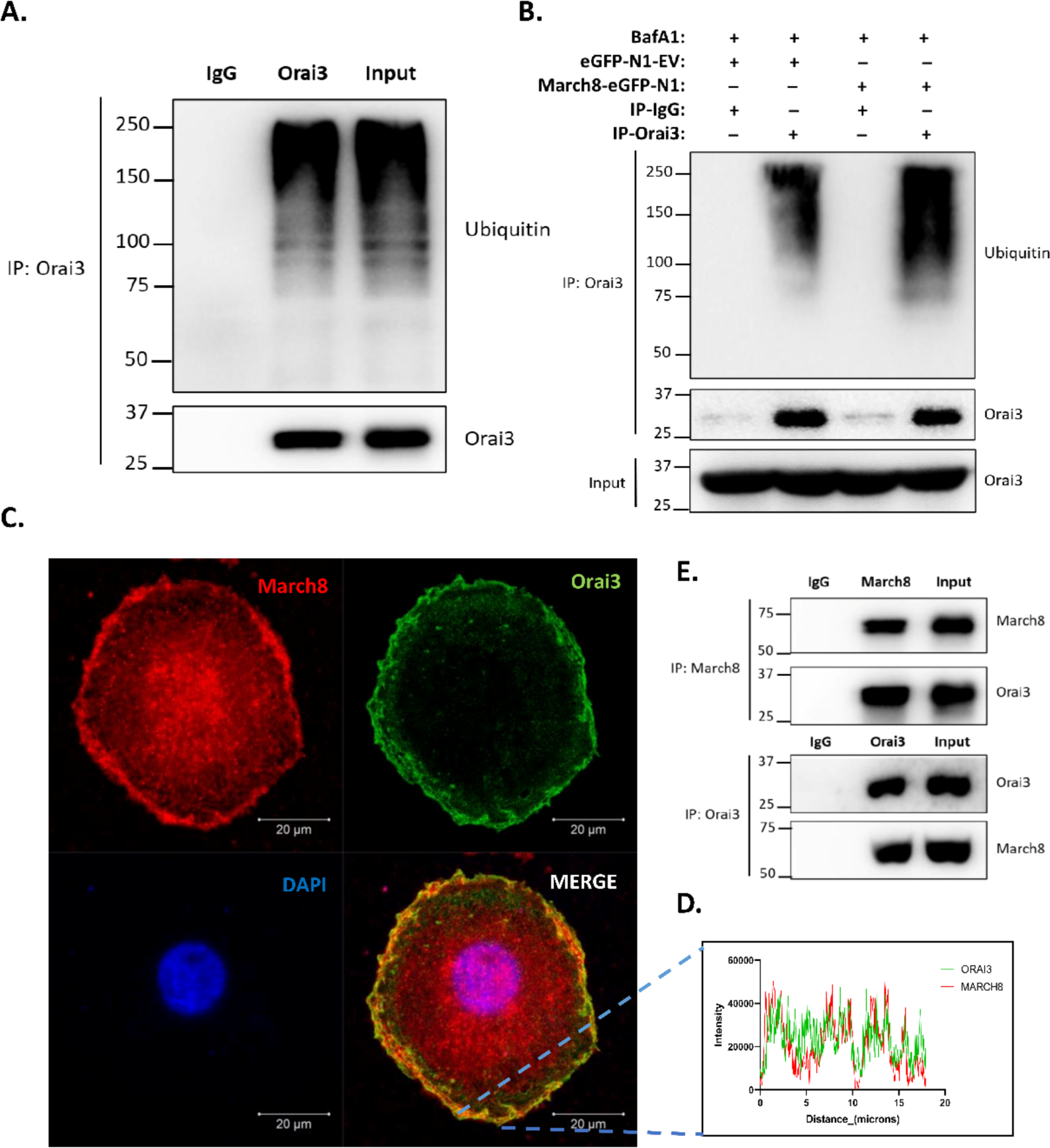
MARCH8 physically associates with Orai3 and induces ubiquitination. **A).** Ubiquitination assay showing ubiquitination of immunoprecipitated Orai3 in bafilomycinA1 (250 nM) treated PANC-1 cells. **B).** Ubiquitination assay showing that overexpression of MARCH8 in bafilomycin (250 nM) treated PANC-1 cells increases ubiquitin levels in immunoprecipitated Orai3 compared to the empty vector control. The input blot shows endogenous Orai3 protein in the PANC-1 cells used for co-immunoprecipitation. **C).** Immunofluorescence assay for colocalization of MARCH8 and Orai3 in PANC-1 cells. **D).** Profile intensity plot of MARCH8 and Orai3 signals in immunofluorescence assay. **E).** Co-immunoprecipitation and reverse Co-immunoprecipitation assay in PANC-1 cells showing interaction of MARCH8 and Orai3. Coimmunoprecipitation assays were performed at least three independent times.

### NFAT2 oppositely regulates MARCH8 expression in non-metastatic v/s invasive and metastatic PC cells

Our investigation into the mechanisms behind Orai3 protein degradation began with the observation that NFAT2 promotes Orai3 degradation in invasive and metastatic pancreatic cancer cells. After identifying that the MARCH8 E3 ubiquitin ligase ubiquitinates Orai3, leading to its lysosomal degradation, one key question still remains: Does NFAT2 regulate MARCH8 levels, thereby driving Orai3 degradation in PC cells? To answer this question, we again utilized the bioinformatic tools to examine human MARCH8 promoter. The bioinformatic analysis with different algorithms and TF position weight matrix combinations showed one NFAT2 binding site on MARCH8 promoter, which is present at -256bp before the transcription start site (Figure 7A). Further, the multi-species alignment of the MARCH8 promoter highlighted that this binding site is conserved across multiple mammalian species (Figure 7A). Additionally, expression analysis of NFAT2 and MARCH8 using the database “GEPIA” showed a positive correlation between NFAT2 and MARCH8 mRNA expression in pancreatic tissue (Figure 7B). To validate the bioinformatic findings, we conducted dual luciferase assays in which the human MARCH8 promoter was cloned into a luciferase reporter vector. These assays were performed in PANC-1 cells with NFAT2 overexpression. We observed an increase in relative luciferase activity upon NFAT2 overexpression in comparison to the empty vector control (Figure 7C). We next carried out Chromatin Immunoprecipitation (ChIP) assays to examine NFAT2 binding on the MARCH8 promoter. We overexpressed either EGFP-C1-NFAT2 or the empty vector control (pEGFP-C1) for ChIP assays. The cross-linked and sonicated DNA was immunoprecipitated and amplified using human MARCH8 promoter-specific primers. ChIP-qPCR analysis revealed a significant enrichment of the MARCH8 promoter with potential NFAT2 binding site compared to mock IP samples (Figure 7D). Collectively, the findings from the ChIP and dual luciferase assays clearly demonstrate that NFAT2 binds to the MARCH8 promoter.

**Figure 7:**
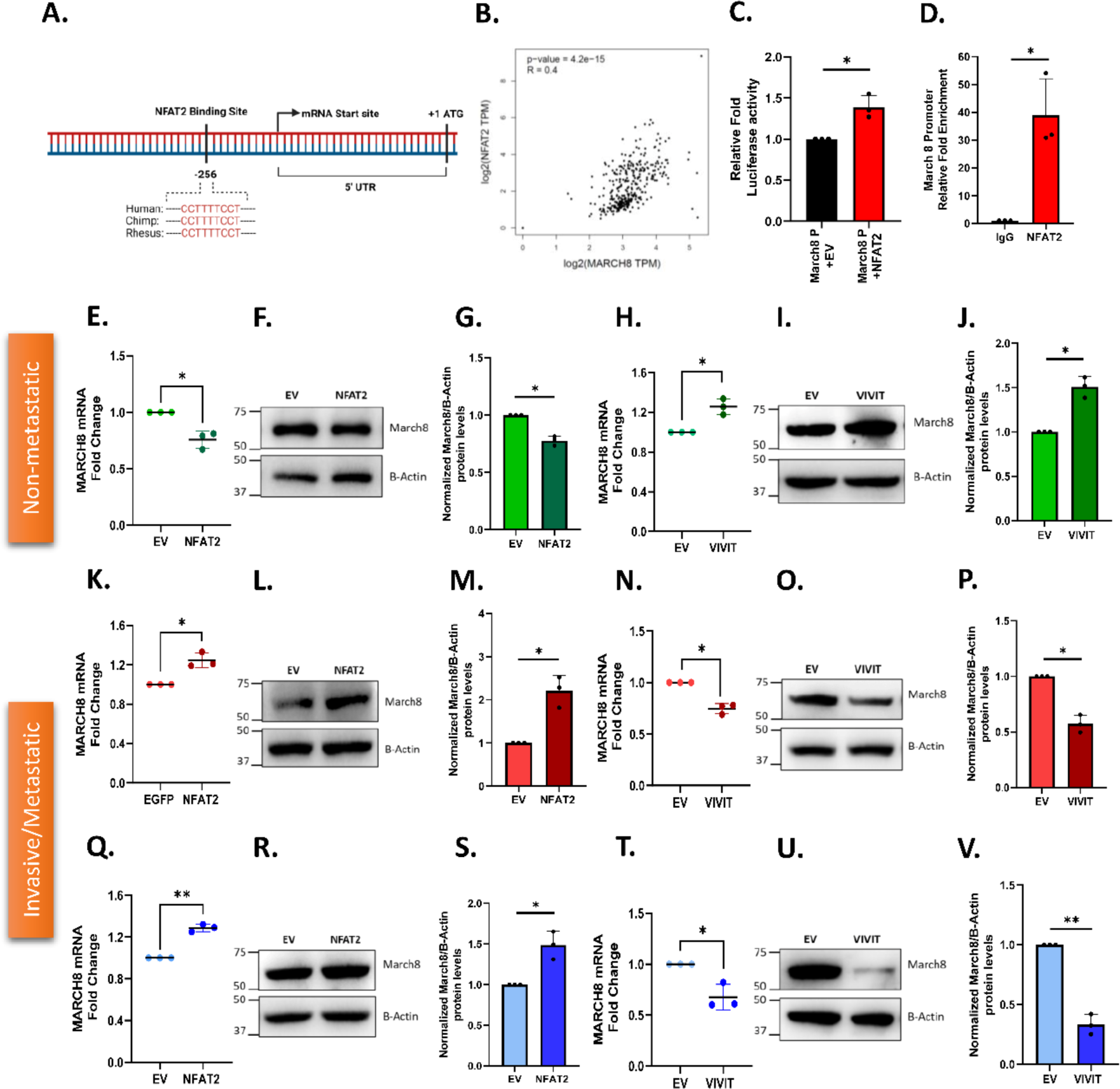
NFAT2 differently regulates MARCH8 expression in non-metastatic v/s metastatic PC cells. **A).** Identification of putative NFAT2 binding sites on the Human MARCH8 promoter using the EPD-Search Motif Tool at p-val cut-off of 0.01 and bioinformatic characterization of conserved NFAT2 binding sites on the cross-species alignment of the MARCH8 promoter using the ContraV3 transcription factor binding analysis tool. **B).** NFAT2 and MARCH8 mRNA expression analysis in GEPIA showing a positive correlation of NFAT2 and MARCH8 in pancreatic tissues. **C).** Normalized Luciferase activity of MARCH8 promoter in PANC-1 cells upon NFAT2 overexpression for 48 hrs. (N=3). **D).** ChIP-qPCR Analysis for relative fold enrichment in NFAT2 immunoprecipitated DNA samples showing higher enrichment of MARCH8 promoter region compared to IgG mock IP (N=3). **E).** qRT-PCR analysis showing decrease in MARCH8 mRNA levels upon NFAT2 overexpression in MiaPaCa-2 compared to control (N=3). **F).** Representative western blots for decrease in MARCH8 protein levels due to NFAT2 overexpression in MiaPaCa-2 cells compared to control. **G).** Densitometric quantitation of MARCH8 protein levels in NFAT2 overexpressed MiaPaCa-2 compared to control (N=3). **H).** qRT-PCR analysis showing increase in MARCH8 mRNA levels upon VIVIT transfection in MiaPaCa-2 compared to control (N=3). **I).** Representative western blots for increase in MARCH8 protein levels due to VIVIT transfection in MiaPaCa-2 cells compared to control. **J).** Densitometric quantitation of MARCH8 protein levels in VIVIT transfected MiaPaCa-2 compared to control (N=3). **K).** qRT-PCR analysis showing increase in MARCH8 mRNA levels upon NFAT2 overexpression in PANC-1 compared to control (N=3). **L).** Representative western blots for increase in MARCH8 protein levels due to NFAT2 overexpression in PANC-1 cells compared to control. **M).** Densitometric quantitation of MARCH8 protein levels in NFAT2 overexpressed PANC-1 compared to control (N=3). **N).** qRT-PCR analysis showing decrease in MARCH8 mRNA levels upon VIVIT transfection in PANC-1 compared to control (N=3). **O).** Representative western blots for decrease in MARCH8 protein levels due to VIVIT transfection in PANC-1 cells compared to control. **P).** Densitometric quantitation of MARCH8 protein levels in VIVIT transfected PANC-1 compared to control (N=3). **Q).** qRT-PCR analysis showing increase in MARCH8 mRNA levels upon NFAT2 overexpression in CFPAC-1 compared to control (N=3). **R).** Representative western blots for increase in MARCH8 protein levels due to NFAT2 overexpression in CFPAC-1 cells compared to control. **S).** Densitometric quantitation of MARCH8 protein levels in NFAT2 overexpressed CFPAC-1 compared to control (N=3). **T).** qRT-PCR analysis showing decrease in MARCH8 mRNA levels upon VIVIT transfection in CFPAC-1 compared to control (N=3). **U).** Representative western blots for decrease in MARCH8 protein levels due to VIVIT transfection in CFPAC-1 cells compared to control. **V).** Densitometric quantitation of MARCH8 protein levels in VIVIT transfected CFPAC-1 compared to control (N=3). Data presented are mean ± S.E.M. For statistical analysis, one sample t-test was performed for panels C, D, E, G, H, J, K, M, N, P, Q, S, T and V using GraphPad Prism software. Here, * *p* <0.05; and ** *p* < 0.01.

To delineate the role of NFAT2 in regulating MARCH8 in PC cells, we performed NFAT2 overexpression and VIVIT transfection studies in MiaPaCa-2, PANC-1, and CFPAC-1 cell lines. qRT-PCR analysis showed that overexpression of NFAT2 in MiaPaCa-2 (non-metastatic) decreased MARCH8 mRNA levels compared to the control (Figure 7E). While NFAT2 overexpression increased MARCH8 mRNA levels in invasive PANC-1 (Figure 7K) and metastatic CFPAC-1 (Figure 7Q). Likewise, western blot analysis showed the same trend; NFAT2 overexpression decreases Orai3 protein levels in MiaPaCa-2 (Figure 7F**, G**) while it increased Orai3 protein expression in PANC-1 (Figure 7L**, M**) and CFPAC-1 cells (Figure 7R**, S**). Next, we performed qRT-PCR analysis upon VIVIT transfection in MiaPaCa-2 and observed an increase in MARCH8 mRNA levels compared to the control (Figure 7H). However, VIVIT transfection decreased MARCH8 mRNA levels in PANC-1 (Figure 7N) and CFPAC-1 (Figure 7T). Similarly, western blot analysis demonstrated the same trend i.e. VIVIT transfection increases Orai3 protein levels in MiaPaCa-2 (Figure 7I**, J**) while it decreases Orai3 protein levels in PANC-1 (Figure 7O**, P**) and CFPAC-1 cells (Figure 7U**, V**). Taken together, these data establish NFAT2 as a repressor of MARCH8 transcription in non-metastatic cells and as an activator in invasive and metastatic PC cells.

### Epigenetic landscape of MARCH8 promoter dictates NFAT2-driven Orai3 regulation in non-metastatic v/s metastatic PC cells

If a transcription factor regulates its target gene differently across distinct cells, it could be because of differences in the epigenetic modifications in promoter region of the target gene (Gibney & Nolan, 2010). One of the most common epigenetic modifications is DNA methylation, and changes in the methylation profile especially in the promoter region leads to differences in gene expression (Lakshminarasimhan & Liang, 2016). Therefore, we investigated the potential contribution of DNA methylation in NFAT2’s dichotomous role in regulating MARCH8 transcription. To check whether the MARCH8 gene promoter is methylated, we used the DNMIVD database (Ding et al., 2020) and found that the MARCH8 promoter is hypermethylated in pancreatic tumor samples compared to normal pancreatic samples (Figure 8A). Consequently, the expression of MARCH8 is lesser in pancreatic tumor samples compared to normal pancreatic samples (Figure 8B). Hence, the expression profile of MARCH8 and the methylation status of the MARCH8 promoter are inversely correlated (Figure 8C). Further, we looked for CpG islands in MARCH8 promoter using Methprimer (Li & Dahiya, 2002) and UCSC genome browser (Karolchik et al., 2009) and found one CpG island extending from -108 bp to +483 bp from the transcription start site (TSS) (Figure 8D). Collectively, bioinformatic analysis suggests that the MARCH8 promoter is hypermethylated in pancreatic cancer and regulates its expression.

**Figure 8:**
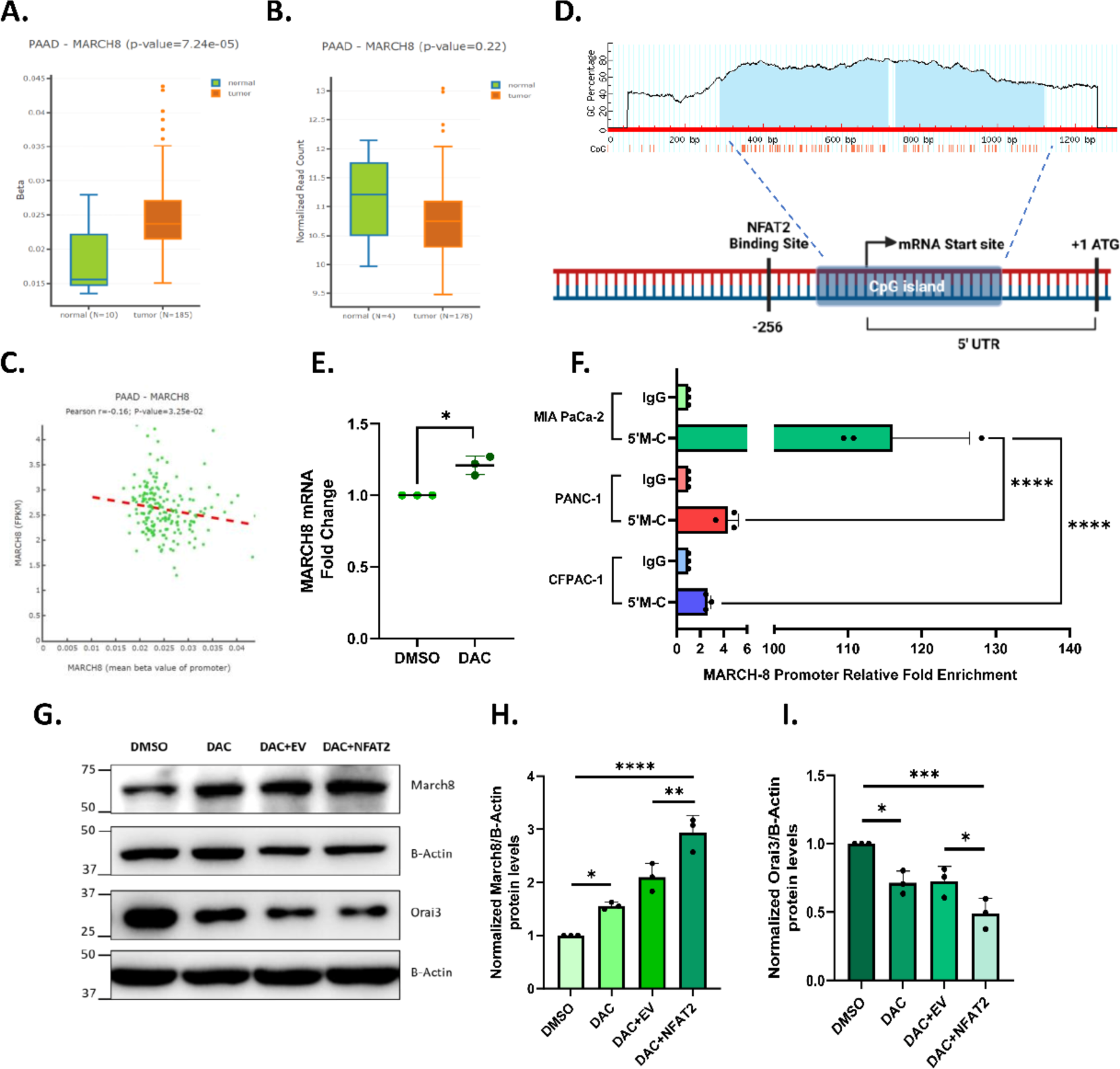
Epigenetic landscape of MARCH8 Promoter dictates NFAT2 driven regulation in non-metastatic v/s metastatic PC cells. **A).** DNA methylation levels of the MARCH8 promoter region between normal tissues and pancreatic adenocarcinoma (PAAD) tissues analysed by the DNMIVD database. **B).** MARCH8 expression levels in normal pancreatic tissues and pancreatic adenocarcinoma (PAAD) tissues analysed by the DNMIVD database. **C).** Correlation between MARCH8 expression and DNA methylation level of MARCH8 promoter in PAAD tissues. **D).** Schematic showing location of CpG island in MARCH8 promoter. **E).** qRT-PCR analysis showing increase in MARCH8 mRNA levels upon decitabine (5µM) treatment in MiaPaCa-2 compared to control (N=3). **F).** MeDIP qPCR Analysis for relative fold enrichment in 5-methylcytosine immunoprecipitated DNA samples showing higher enrichment of MARCH8 promoter region in MiaPaCa-2 compared to PANC-1 and CFPAC-1 (N=3). **G).** Representative western blots for increase in MARCH8 levels and decrease in Orai3 levels upon NFAT2 overexpression in decitabine treated MiaPaCa-2 cells. **H** & **I)**. Densitometric analysis of MARCH8 and Orai3 levels upon NFAT2 overexpression in decitabine treated MiaPaCa-2 cells. Data presented are mean ± S.E.M. For statistical analysis, one sample t-test was performed for panels E and F while ordinary one-way ANOVA was performed followed by Tukey’s multiple comparison test for panels H and I using GraphPad Prism software. Here, * *p* <0.05; *** *p* < 0.001; and **** *p* < 0.0001.

To corroborate the lead from the bioinformatic analysis, we treated MiaPaCa-2 cells with DNA methylation inhibitor, decitabine. Decitabine (5-Aza-2′-Deoxycytidine) is a deoxycytidine analog that integrates into DNA and inhibits DNA methyltransferase activity, thereby preventing DNA methylation (Jones & Taylor, 1980). qRT-PCR analysis shows that decitabine treatment in MiaPaCa-2 for 72hr significantly increases MARCH8 mRNA levels compared to the vehicle control (Figure 8E). We next examined the methylation status of the MARCH8 promoter in non-metastatic, invasive, and metastatic cells. We performed MeDIP (Methyl DNA immunoprecipitation) using antibodies against 5-methylcytosine. We observed that the methylated MARCH8 promoter was highly enriched in MiaPaCa-2 compared to PANC-1 and CFPAC-1, suggesting that the MARCH8 promoter is hypermethylated in non-metastatic cells compared to invasive and metastatic cells (Figure 8F). Finally, to investigate whether DNA methylation prevents NFAT2 from regulating MARCH8 in non-metastatic cells, we overexpressed NFAT2 in decitabine-treated MiaPaCa-2 cells. We found that MARCH8 protein levels increased compared to both vehicle control and empty vector control plus decitabine-treated cells (Figure 8G**, H**). Consequently, due to an increase in MARCH8 levels, Orai3 protein levels decreased in NFAT2 overexpressed decitabine-treated cells compared to both vehicle control and empty vector control plus decitabine-treated cells (Figure 8G**, I**). Taken together, our data demonstrates that the MARCH8 promoter is hypermethylated in non-metastatic cells and thereby NFAT2 does not upregulate MARCH8 in these cells. This in turn prevents NFAT2-induced Orai3 degradation, which is observed in invasive and metastatic cells.

### MARCH8 E3 ubiquitin ligase abrogates pancreatic cancer metastasis

Our study has identified MARCH8 as a critical regulator of Orai3, which drives PC metastasis (Arora et al., 2021). Although MARCH8 E3 ubiquitin ligase has been studied in some cancer types (W. Chen et al., 2021; Fan et al., 2017; Singh et al., 2017; Z. Wang et al., 2022), its role in PC remains largely unappreciated. Therefore, we examined the relationship between elevated MARCH8 expression and the survival time of PC patients using data from the ‘OncoLnc’ database. This analysis revealed that the patients with low MARCH8 expression survived for a shorter duration as compared to patients with high MARCH8 expression (Figure 9A). This highlights that higher MARCH8 levels are associated with better prognosis in PC patients. It further suggests that MARCH8 could play a role in PC progression, and metastasis that eventually leads to mortality. To investigate MARCH8’s role in PC progression and metastasis, we generated stable CFPAC-1 cell lines using lentiviral transduction, expressing either a non-targeting control shRNA (shNT) or an shRNA specifically targeting MARCH8 (shMARCH8). Western blot analysis confirmed a marked reduction in MARCH8 protein levels in CFPAC-1 shMARCH8 cells compared to the shNT control (Figure 9B**, C**). After confirming the knockdown, we performed scratch wound assays to explore the role of MARCH8 in CFPAC-1 cell migration. Cell migration post-scratch was monitored by assessing wound closure temporally. The results showed that CFPAC-1 shMARCH8 cells closed the wound more effectively than shNT cells (Figure 9D), shMARCH8 cells closed 50% of the wound within 24 hours, while shNT cells closed less than 20% (Figure 9E). This demonstrates the involvement of MARCH8 in regulating pancreatic cancer cell migration. Finally, to examine the role of MARCH8 in PC metastasis *in vivo*, we carried out zebrafish xenograft experiments by injecting stable CFPAC-1 cells into the peri-vitelline space (PVS) of 2-day post-fertilization (dpf) zebrafish (Figure 9F) and monitored metastatic events three hours post-injection (Martinez-Lopez et al., 2021; White & Patton, 2023). As shown in Figure 9G-H, zebrafish injected with CFPAC-1 shMARCH8 cells exhibited increased metastatic events compared to those injected with CFPAC-1 shNT cells. Altogether, the *in vitro* and *in vivo* data highlights that MARCH8 acts a negative regulator of PC metastasis.

**Figure 9:**
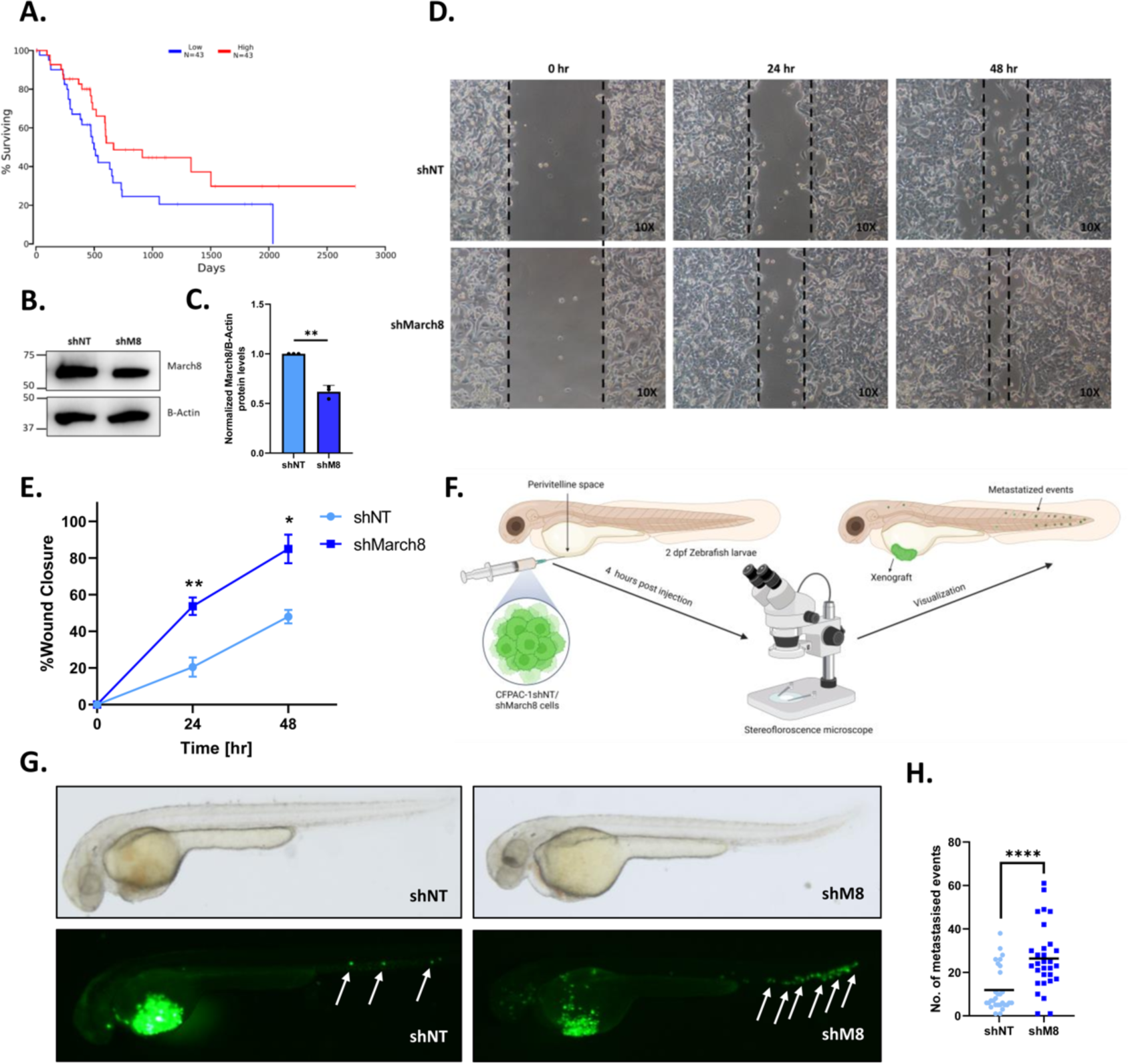
MARCH8 E3 ubiquitin ligase abrogates metastatic potential in pancreatic cancer. **A).** Survival Analysis in pancreatic cancer patients wherein blue trace corresponds to low MARCH8 expression (n=43), and red trace corresponds to high MARCH8 expression (n=43) highlighting that high MARCH8 expression increases patient’s survival time. **B).** Representative western blots for MARCH8 levels in CFPAC1 shNT and shMARCH8 cells. **C).** Densitometric analysis of MARCH8 protein levels in CFPAC-1 shNT control and shMARCH8. **D).** Representative wound healing images of CFPAC-1 shNT and shMARCH8 cells at 0h, 24h and 48h. **E).** Quantitation of % wound closure from CFPAC-1 shNT and shMARCH8 cells over the interval of 48h. **F).** Schematic diagram showing the workflow for zebrafish xenograft injections. **G).** Representative bright-field and florescent images of 2dpf zebrafish injected with GFP-labelled CFPAC-1 shNT and shMARCH8 cells. CFPAC-1 shMARCH8 cells shows increased metastasised events compared to control. Metastasized events are indicated with white arrow marks. **H).** Quantitation of metastasised events in CFPAC-1 shNT and shMARCH8 injected zebrafish (each dot represents data from individual embryos from three independent experiments N=∼30). Data presented are mean ± S.E.M. For statistical analysis, one sample t-test was performed for panel C while unpaired student t-test was performed for panel H using GraphPad Prism software. Here, ** *p* < 0.01; and **** *p* < 0.0001.

To summarize, in this study we have identified that the same transcription factor regulates Orai3 transcription and protein degradation. Firstly, we report that NFAT2 binds to Orai3 promoter and controls its transcription in non-metastatic, invasive and metastatic PC cells. But additionally, NFAT2 activates Orai3 lysosomal degradation system facilitated by MARCH8 E3 ubiquitin ligase in invasive and metastatic PC cells. Through gain of function and loss of function experiments, along with bio-chemical and high-resolution imaging studies, we establish that MARCH8 ubiquitinates and subsequently degrades Orai3. Notably, we show that epigenetic differences in the MARCH8 promoter allow NFAT2 to regulate MARCH8 transcription and promote Orai3 protein degradation in invasive and metastatic cells, while this mechanism is absent in non-metastatic cells. In non-metastatic PC cells, the MARCH8 promoter is hypermethylated and hence hinders NFAT2 guided transcription while in invasive and metastatic PC cells, MARCH8 promoter is relatively hypomethylated. Furthermore, we report a critical role of MARCH8 in PC metastasis, as our *in vitro* and *in vivo* findings suggest that MARCH8 restricts PC metastasis. Collectively, we have identified a dual regulator of Orai3 expression in a context dependent manner (Figure 10). Moreover, we have revealed the therapeutic potential of targeting MARCH8 E3 ubiquitin ligase to curtail PC metastasis.

**Figure 10:**
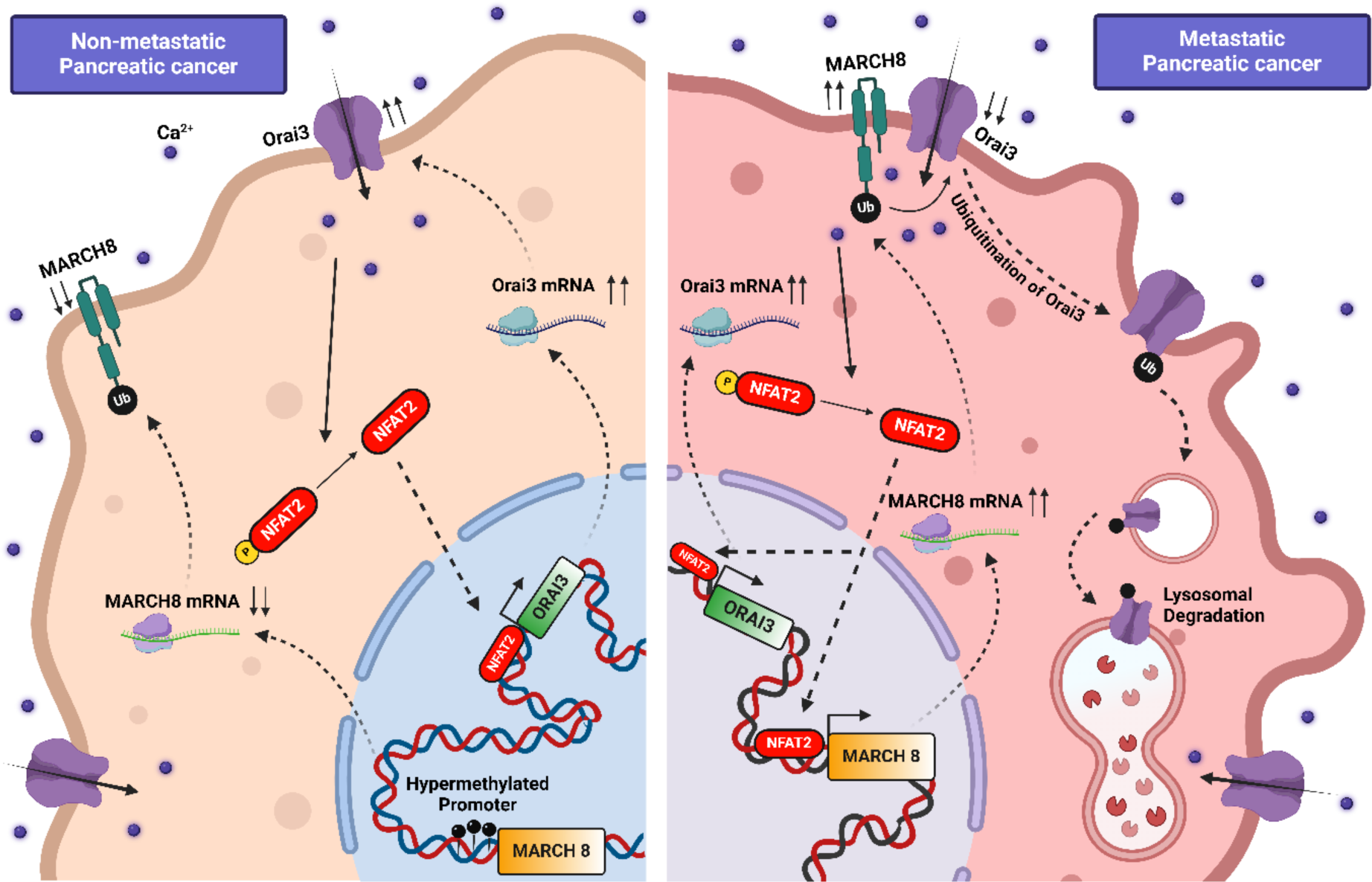
Dual role of NFAT2 in regulating Orai3 transcription and protein degradation. In non-metastatic cells, NFAT2 drives Orai3 transcriptional upregulation whereas in metastatic cells NFAT2 induces Orai3 lysosomal degradation via MARCH8 E3 ubiquitin ligase. The dichotomy in NFAT2’s function in non-metastatic versus metastatic cells is an outcome of differences in the methylation status of MARCH8 promoter in these cells.

## Discussion

Ca^2+^ signaling plays a key role in regulating cancer progression. A previous study from our group revealed a critical role of Orai3 channel in regulating pancreatic cancer development and metastasis *in vivo*. We reported that Orai3 is transcriptionally upregulated in pancreatic tumor samples compared to normal pancreatic samples. The upregulation of Orai3 is well-documented in other types of cancers as well but the molecular players and signaling modules driving Orai3 regulation are still unknown. In this study we discovered that NFAT2 regulates Orai3 expression. NFAT2 belongs to NFATc group of transcription factors. Interestingly, four isoforms of NFATc i.e. NFAT1, 2, 3 and 4 are Ca^2+^ sensitive and are activated upon increase in cytosolic Ca^2+^ levels (Müller & Rao, 2010; Prakriya et al., 2006; Tanwar et al., 2024). Using *in-silico* analysis we found potential binding sites of NFATc on the Orai3 promoter (Figure 1B). Next, we shortlisted NFAT2 as it significantly upregulates Orai3 at both mRNA and protein levels in HEK-293T cells (Figure 1D**, E**) and GEPIA database showed positive correlation of NFAT2 and Orai3 mRNA in pancreatic tissues (Figure 1G).

Interestingly, the NFAT2 overexpression and NFAT inhibition studies in pancreatic cancer cell lines showed a surprising dichotomy in regulation of Orai3 by NFAT2 in non-metastatic v/s invasive and metastatic cells We observed that in invasive and metastatic cells like PANC-1 and CFPAC-1, NFAT2 acts as bimodal regulator to maintain Orai3 levels as NFAT2 overexpression leads to decrease in Orai3 at protein level (Figure 2F**, G, K, L**). This negative feedback loop in invasive and metastatic cells could be because invasive and metastatic cells have higher endogenous Orai3 levels compared to non-metastatic cells (Arora et al., 2021). A further increase in Orai3 levels by NFAT2 would allow excess amount of Ca^2+^ entering into the cell which could lead to apoptosis. While in non-metastatic cells like MiaPaCa-2, NFAT2 transcriptionally upregulates Orai3, leading to increase in Orai3 at protein levels (Figure 2A, Therefore, our data demonstrates that a Ca^2+^ sensitive transcription factor induces transcription of a Ca^2+^ channel thereby indicating presence of a positive feedback loop in these cells. However, further studies are required to decipher the source of rise in cytosolic Ca^2+^ and Ca^2+^ handling toolkit involved in increasing cytosolic Ca^2+^ levels for activating NFAT2.

Another interesting observation from our study is that inhibition of NFAT activity by VIVIT leads to transcriptional upregulation of Orai3 in non-metastatic, invasive and metastatic cells (Figure 3A**, F, K**). We performed luciferase assay in VIVIT transfected PANC1 cells and observed increase in relative luciferase activity in VIVIT transfected cells compared to empty vector control. This indicates that inhibition of NFAT’s activity by VIVIT peptide activates or allows other endogenous transcription factors to bind on Orai3 promoter and upregulate Orai3 transcription. Nonetheless, our data elegantly demonstrates that NFAT is a critical regulator of both Orai3 transcription and degradation.

It is important to highlight that just like transcriptional regulation, Orai3’s protein degradation process is not appreciated yet. The bimodal regulation of Orai3 by NFAT2 in invasive and metastatic pancreatic cancer cells led to characterization of Orai3 protein degradation machinery. Using lysosomal and proteasomal degradation inhibitors, we observed that NFAT2 induces lysosomal degradation of Orai3 in both invasive and metastatic pancreatic cancer cells (Figure 3A**, B, C, D**). Orai3 is a plasma membrane (PM) protein, and most PM proteins are degraded in lysosomes, although the mechanism and the players involved in the degradation process are different. Typically, the PM target proteins are ubiquitinated by E3 ubiquitin ligases and are then delivered to the late endosomes/multivesicular body (MVB) via endocytosis. The endosomes containing ubiquitinated PM proteins eventually fuse with lysosomes and are finally degraded by lysosomal proteases (Zhao et al., 2022). Our unbiased and robust bioinformatics narrowed down to MARCH8 E3 ubiquitin ligase as a potential regulator of Orai3 degradation (Figure 4E). Indeed, MARCH8 overexpression and knockdown experiments show that MARCH8 induces Orai3 degradation in non-metastatic, invasive, and metastatic cells (Figure 4 **& 5**). Mechanistically, MARCH8 ubiquitinates Orai3 and induces lysosomal degradation (Figure 6B). Further, by performing co-immunoprecipitation and immunofluorescence assay, we showed that MARCH8 and Orai3 physically interact (Figure 6C**, D, E**). This data established that MARCH8 E3 ubiquitin ligase physically interacts with Orai3 at plasma membrane and induces lysosomal degradation.

Next, we investigated whether NFAT2 transcriptionally regulates MARCH8 and thereby induces Orai3 protein degradation. To examine this, we did *in-silico* analysis and found that NFAT2 has one potential binding site on human MARCH8 promoter, which is conserved across mammals (Figure 7A). Further, NFAT2 overexpression and VIVIT transfection studies in non-metastatic, invasive and metastatic PC cells establish that NFAT2 acts as a repressor in non-metastatic cells while acting as an activator in invasive and metastatic cells (Figure 7). This clearly explains that the NFAT2 induced Orai3 protein degradation in invasive and metastatic cells is due to increase in MARCH8 expression. Similarly, decrease in Orai3 protein levels in non-metastatic cells upon NFAT inhibition is due to MARCH8 upregulation. The transcriptional regulation of MARCH8 by NFAT2 presents an intriguing area for exploration beyond cancer biology. Both NFAT2 and MARCH8 are prominent players in human immunology. NFAT2 regulates the transcription of critical immune receptors such as ICOS, PD-1, CXCR5, CD40L, and chemokines like IL-4 and IL-21 (Vaeth & Feske, 2018). While MARCH8, an E3 ubiquitin ligase, targets MHC-II and CD80 receptors and plays a crucial role in antiviral defense (Lin et al., 2019b). Therefore, investigating the relationship between NFAT2 and MARCH8 in immune cells would be beneficial for advancing our understanding of cancer immunotherapy, immune disorders and infectious disease biology.

The dual role of NFAT2 as a repressor and activator of MARCH8 in pancreatic cancer cells depending on the cell type points out that there could be epigenetic differences in the MARCH8 promoter region. Indeed, using *in-silico* analysis we found that indeed the MARCH8 promoter is epigenetically modified, more specifically shows DNA methylation. DNA methylation is one of the most common epigenetic modifications in the genome and generally DNA methylation suppresses gene activation (Nishiyama & Nakanishi, 2021). Interestingly, we found that MARCH8 promoter is hypermethylated in pancreatic tumor samples compared to normal pancreatic samples (Figure 8A) and consequently, the expression of MARCH8 is lower in tumor samples compared to normal pancreatic samples (Figure 8B), showing negative correlation (Figure 8D). Using MeDIP assay, we show that there are differences in DNA methylation levels of MARCH8 promoter in non-metastatic, invasive and metastatic cell lines. Our results demonstrate that the MARCH8 promoter is hypermethylated in non-metastatic PC cells (Figure 8F). To examine if this methylation hinders NFAT2 from regulating MARCH8 transcription in non-metastatic cells, we overexpressed NFAT2 in MiaPaCa-2 cells treated with DNA methylase inhibitor (decitabine). Western blot analysis revealed that NFAT2 overexpression in decitabine-treated cells led to an increase in MARCH8 levels compared to other control conditions (Figure 8G**, H, I**). In summary, the MARCH8 promoter is hypermethylated in non-metastatic PC cells, preventing NFAT2 from promoting MARCH8-mediated Orai3 protein degradation in these cells.

Interestingly, MARCH8 regulates cancer progression both positively and negatively depending upon the type of cancer (C. Chen et al., 2023; W. Chen et al., 2021; Xu et al., 2023; Ying et al., 2024). MARCH8’s role in regulating tumorigenesis depends on its degradation targets. For instance, in breast cancer cells, MARCH8 ubiquitinates STAT3 and CD44, suppressing tumor metastasis (W. Chen et al., 2021) while in hepatocellular carcinoma, MARCH8 induces degradation of PTEN, promoting malignancy (Xu et al., 2023). However, role of MARCH8 in pancreatic cancer biology remains poorly appreciated. Here, our functional studies with stable knockdown of MARCH8 signifies its role in pancreatic cancer migration (Figure 9D**, E**). More importantly, our *in vivo* zebrafish xenograft experiments show that stable knockdown of MARCH8 increases metastatic potential of the pancreatic cancer cells (Figure 9G**, H**). It is important to note that we have earlier reported the Orai3 positively drives PC progression and metastasis (Arora et al., 2021). Therefore, MARCH8 controls PC metastasis at least partially by modulating Orai3 expression and activity.

To summarize, we reveal that NFAT2 regulates both the transcription and lysosomal degradation of the Orai3 oncochannel in a context-dependent manner. In non-metastatic cancer cells, NFAT2 drives Orai3 transcription, increasing its levels. In contrast, in invasive and metastatic cells, NFAT2 promotes Orai3 degradation by upregulating the MARCH8 E3 ubiquitin ligase. Biochemical assays and super-resolution microscopy confirm that MARCH8 physically interacts with Orai3, leading to its degradation. This regulatory shift is linked to the epigenetic differences, as the MARCH8 promoter is highly methylated in non-metastatic cells, preventing NFAT2 from inducing Orai3 degradation. The dual regulation of Orai3 by same transcription factor NFAT2 underscores a complex pathway in pancreatic cancer cells that controls disease progression and clinical outcomes.

## Methodology

### Materials and Reagents

Calcium Imaging reagents like thapsigargin and 2-APB were purchased from Sigma. FURA-2AM was obtained from Invitrogen. MG-132, Bafilomycin-A1 and Decitabine (5-Aza-2′-Deoxycytidine) were procured from Sigma. MARCH8-eGFP-N1 WT and MARCH8-eGFP-N1 CS plasmids were gifted by Hideaki Fujita (Fujita et al., 2013). Orai3-eYFP plasmid was a gift from Mohamed Trebak.

### Cell Culture

Human embryonic kidney cell line HEK-293T and human pancreatic cancer cell lines MiaPaCa-2, PANC-1 and CFPAC-1 were acquired from ATCC. CFPAC-1 were cultured in RPMI-1640 supplemented with L-Glutamine. MiaPaCa-2 were cultured in DMEM medium with 2.5% Horse serum and 10% fetal bovine serum (FBS). HEK 293-T and PANC-1 were cultured in DMEM added with 10% FBS.

### qRT-PCR Analysis

For the extraction of mRNA, the cell pellets were processed with Qiagen RNeasy kit (Catalog #74106). cDNA was made from the extracted mRNA via high-capacity cDNA reverse transcription kit from Thermo Fisher Scientific (Catalog #4368814). Real-time PCR were performed using SYBR green in Applied Biosystems Quant Studio 6 Flex. The analysis was done with Quant Studio RT-PCR software version 1.3. The mRNA expression of NFAT2, Orai3 and MARCH8 in experimental conditions were normalized to GAPDH (Housekeeping gene). The primer sequences used are as follows:

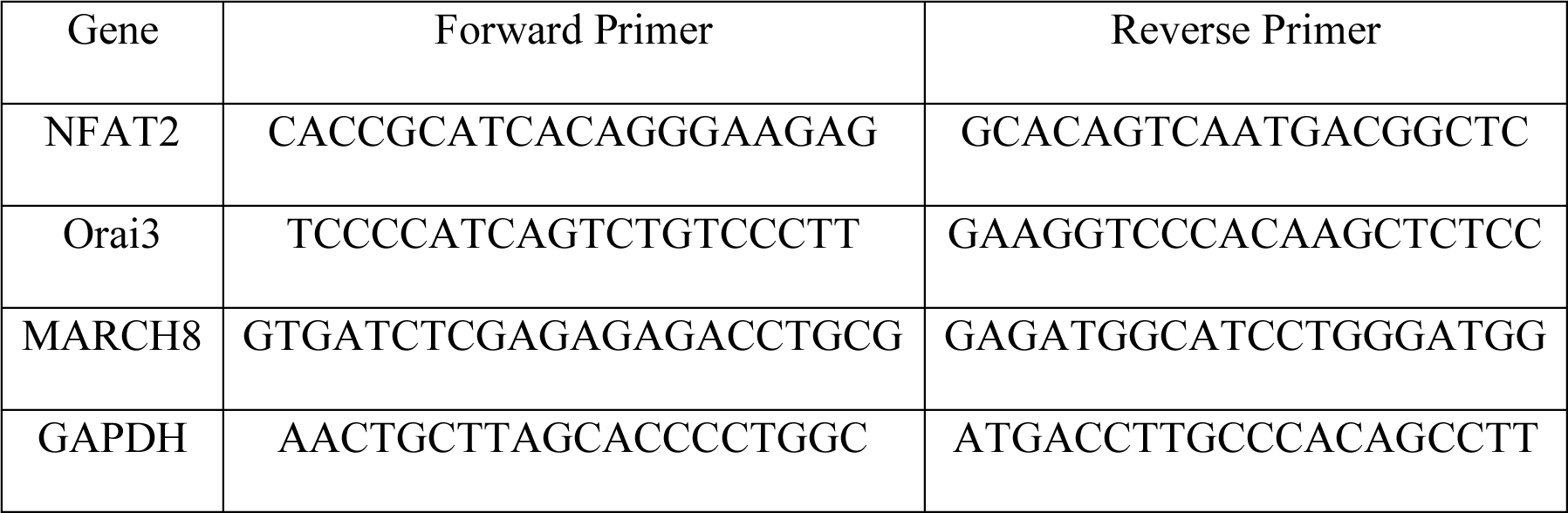

### Western Blotting

The total protein from the cell pellets of experimental conditions were extracted using NP40 lysis buffer with protease inhibitor cocktail. The quantification of total protein concentration was done via BCA assay (Thermo Fisher Scientific, Catalog #23227). The extracted protein samples were separated by 10% SDS-PAGE and electro transferred to PVDF membrane (Merck Millipore, Catalog #IPVH00010). Blocking of the membranes were done with 5% skimmed milk in Tris-buffered saline with Tween 20 (TBST) at room temperature for 2 h, and then incubated overnight at 4°C with specific primary antibody in TBST, like rabbit anti-Orai3 (1:500, Abcam), rabbit anti-MARCH8 (1:1000, Sigma) and mouse anti-Ubiquitin (1:1000, Santa Cruz). After the overnight incubation, the membranes are incubated in horseradish peroxidase conjugated donkey anti-rabbit secondary antibody (1:10000, Sigma) and horseradish peroxidase conjugated horse anti-mouse secondary antibody (1:1000, Cell Signalling Technologies) for 2 h at room temperature, Finally, the membranes were detected using the Immobilon Forte Western HRP Substrate (Merck Millipore, Catalog #WBLUF0500). β-tubulin (1:5000, Abcam) and β-Actin (1:10000, Abcam) were used as loading controls. Densitometric analysis of the blots were achieved using ImageJ software. The list of antibodies used for western blotting are listed below:

| Antibody | Company | Catalog no. |
| --- | --- | --- |
| Orai3 | Abcam | ab254260 |
| MARCH8 | Sigma Aldrich | HPA014597-100UL |
| Ubiquitin | Santa Cruz | sc-271289 |
| β-tubulin | Abcam | ab21058 |
| β-Actin | Abcam | ab8226 |

### Cloning of Orai3 and MARCH8 Promoter

The human Orai3 promoter and MARCH8 promoter was attained from Eukaryotic Promoter Database (EPD) and the sequence was analysed using NCBI-BLAST to find mRNA start site and the first codon. Primers were designed to amplify 1055 bp region (-1024 to +31 w.r.t start codon) of the Orai3 core promoter and 1042 bp region (-989 to +50 w.r.t start codon) of the MARCH8 core promoter respectively. PCR amplification of the Orai3 promoter and MARCH8 promoter was done using Phusion High Fidelity Polymerase (F503, Thermo Fisher Scientific) and the PCR amplicon was cloned into pGL4.23 luciferase reporter vector (Promega) at the KpnI/HindIII sites for Orai3 promoter and KpnI/NheI sites for MARCH8 promoter respectively. The clones were verified through restriction digestion. The primers used for cloning are listed below.

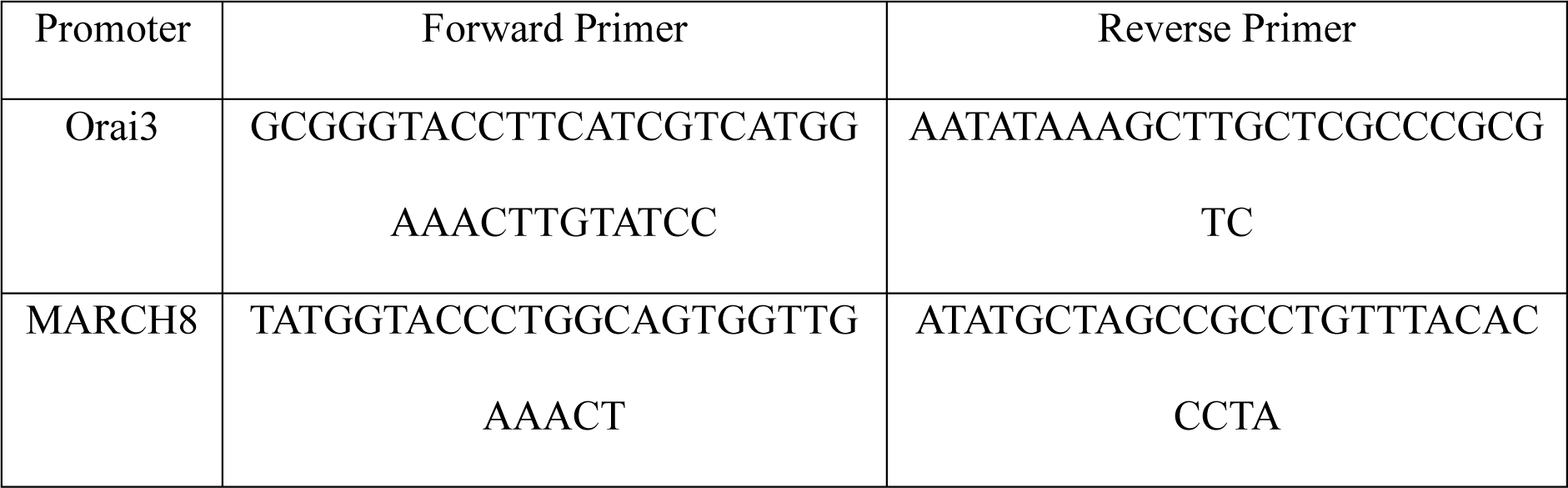

### *In vitro* dual luciferase assay

Before transfection, PANC-1 cells were seeded at a density of 0.5 × 10^5^ cells/well in 24-well plates. Cells were transfected with Orai3P and eGFPC1-NFAT2, and MARCH8P and eGFPC1-NFAT2 as indicated, using jetPRIME transfection reagent (114-01, Polyplus) as per the manufacturer’s protocol. For transfection normalization, renilla luciferase plasmid was used. The cells were assayed for luciferase activity using the dual luciferase assay kit (E1910, Promega) as per manufacturer’s protocol 48 h post transfection. Data is representative of three biological replicates. EGFPC1-huNFATc1EE-WT was a gift from Jerry Crabtree (Addgene plasmid # 24219).

### Chromatin Immunoprecipitation (ChIP)

PANC-1 cells transfected with EGFPC1-huNFATc1EE-WT overexpression plasmid or pEGFPC1-empty vector and were trypsinized after 48 h. The cell count was done and 25 million cells aliquot was resuspended in 10 ml 1× PBS; The aliquots were fixed with 1% formaldehyde for 3 min at room temperature then quenched with 1M Glycine. After quenching, the cells were washed once with cold 1× PBS and pelleted using centrifugation at 1500 rpm for 5 min at 4°C. To shear the chromatin, cell pellets were resuspended in 10 ml of Rinse Buffer 1 (50 mM Hepes pH 8, 140 mM NaCl, 1 mM EDTA, 10% glycerol, 0.5% NP-40, 0.25% Triton X-100) and incubated for 20 min on ice. Cells were next resuspended in Rinse Buffer 2 (10 mM Tris pH 8, 1 mM EDTA, 0.5 mM EGTA, 200 mM NaCl) and incubated on ice for 20 min. Finally, cells were resuspended in Shearing buffer (0.1% SDS, 1 mM EDTA, 10 mM Tris pH 8). For sonication, five million cells were resuspended in 200 µl of shearing buffer and sonicated using Bioruptor Pico with 30s ON and 30s OFF for 20 cycles at 4°C. Sonicated chromatin was centrifuged at 16000 rpm for 10 min to remove debris. DNA concentration of the sonicated chromatin was determined and 25 µg of sonicated chromatin was used for immunoprecipitation using 2 µg GFP antibody or IgG control. The protein A agarose beads (Merck Millipore, Catalog #16-125) used for immunoprecipitation were first equilibrated with glycerol IP buffer and blocking was done using 75 ng/µl Herring sperm DNA and 0.1 µg/µl bovine serum albumin. Post immunoprecipitation, the beads were washed twice each with low salt buffer (0.1% SDS, 1% Triton X100, 2 mM EDTA, 20 mM Tris–Cl pH 8, 150 mM NaCl), high salt buffer (0.1% SDS, 1% Triton X100, 2 mM EDTA, 20 mM Tris– Cl pH 8, 500 mM NaCl), LiCl buffer (0.25 M LiCl, 1% NP-40, 1% sodium deoxycholate, 1 mM EDTA, 10 mM Tris–Cl pH 8) and TE buffer. DNA was eluted from the beads using Elution Buffer (1% SDS, 100 mM NaHCO_3_) for 30 min at 30°C and decrosslinking was performed at 80°C for 1h. Purification of DNA using phenol-chloroform extraction was performed and used for ChIP-qPCR using primers specific to human Orai3 core promoter and MARCH8 core promoter. The primers used for ChIP analysis are as follows:

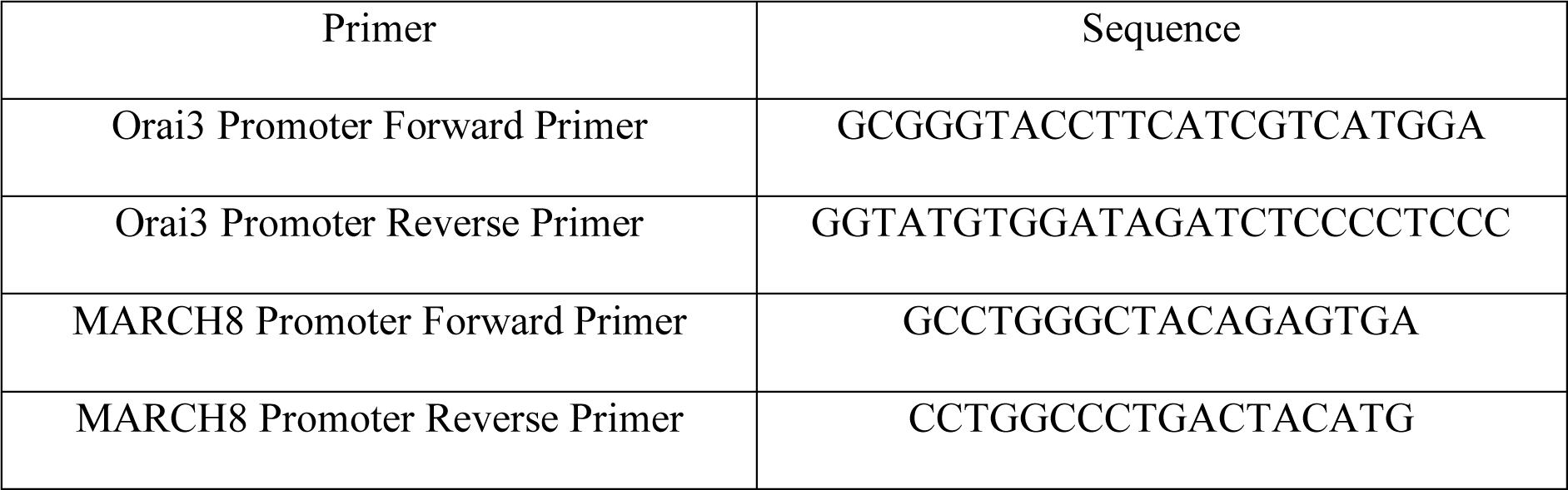

### Calcium Imaging

Calcium (Ca^2+^) imaging was performed as reported earlier (Arora et al., 2021). Briefly, cells were cultured on confocal dishes for performing Ca^2+^ imaging. Cells are incubated at 37°C for 30 min in a culture medium containing 4 µM fura-2 AM. After incubation, cells were washed 3 times and bathed in HEPES-buffered saline solution (140 mM NaCl, 1.13 mM MgCl_2_, 4.7 mM KCl, 2 mM CaCl_2_, 10 mM D-glucose, and 10 mM HEPES; pH 7.4) for 5 min before Ca^2+^ measurements were made. A digital fluorescence imaging system (Nikon Eclipse Ti2 microscope coupled with CoolLED pE-340 Fura light source and a high speed PCO camera) was used, and fluorescence images of several cells were recorded and analysed. Fura-2AM was excited alternately at 340 and 380 nm, and the emission signal was captured at 510 nm. Figures showing Ca^2+^ traces are an average from several cells (the number of cells is denoted as “N” on each trace) attached on a single imaging dish. Each experiment was performed at least 3–4 times and the final data are plotted in the form of bar graphs. The data shown in a particular Ca^2+^ imaging trace originates from multiple cells on a single imaging dish. The exact number of cells and traces for each condition are specified in the respective figure.

### siRNA-based transient transfections

siRNA transfections were done in MiaPaCa-2, PANC-1 and CFPAC-1 cells seeded in 6 well plates. 100nM siNT (Human) and siMARCH8 (Human) were transfected in cells using jetPRIME transfection reagent (114-01, Polyplus) as per the manufacturer’s protocol. Cells were harvested post 48 h of transfection for mRNA and protein expression changes. The siRNAs (smartpool of four individual siRNAs targeting gene of interest) were procured from Dharmacon. The catalog number and target sequence of siRNAs used in the study are as follows:

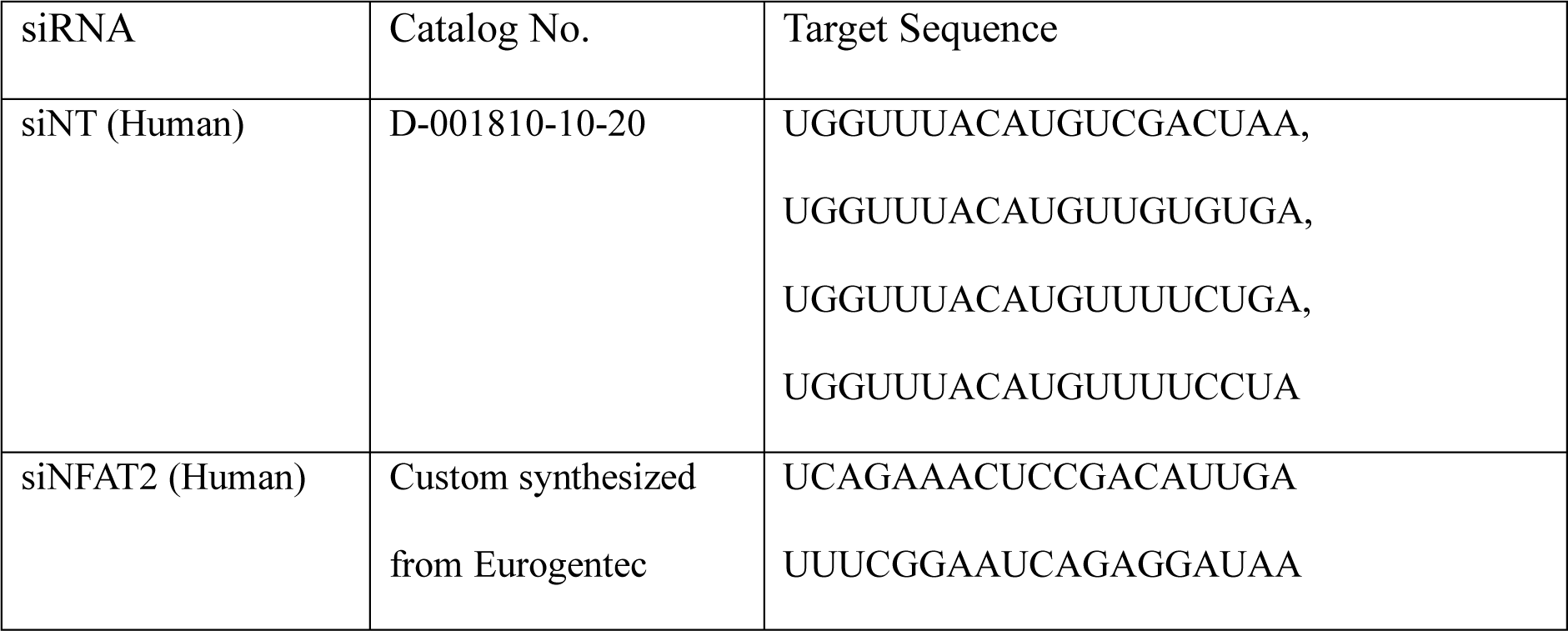

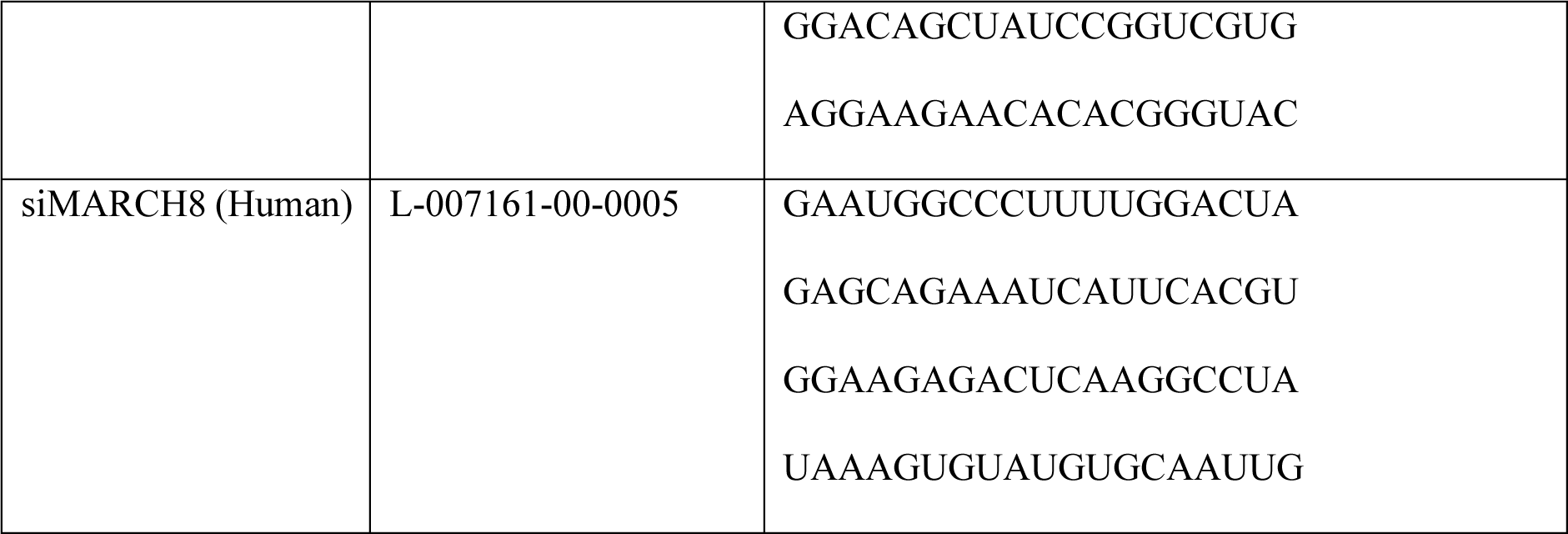

### Ubiquitination Assay

For the ubiquitination assay, PANC-1 cells were cultured in T-75 flasks and after reaching confluency, the cells were collected and lysed with glycerol IP buffer. The protein content in the lysate was calculated using BCA assay and from the cell lysate, 1 mg of protein was used to perform immunoprecipitation using antibody against Orai3. Next day, the incubated lysates were precipitated by adding activated protein A agarose beads and incubating them overnight at 4°C. Before immunoprecipitation, the protein A agarose beads were equilibrated with glycerol IP buffer and blocked with 0.1 µg/µl bovine serum albumin. Precipitation was also performed using IgG (Cell Signaling Technologies) as an isotype control, raised in the same host. The resulting antigen-antibody complexes were processed in NP-40 lysis buffer for immunoblotting to check the levels of ubiquitin and Orai3. To study the effect of MARCH8 on Orai3 ubiquitination, PANC-1 cells were transfected with MARCH8-eGFP-N1 WT or pEGFPN1-empty vector using jetPRIME transfection reagent (114-01, Polyplus) as per the manufacturer’s protocol for 48 hr. Before termination, the cells were treated with bafilomycin (250 nM) for 6 hr. Further the cells were processed as mentioned above.

### Co-Immunoprecipitation

For Co-immunoprecipitation, PANC-1 cells were lysed in glycerol IP buffer for 20 minutes on incubation on ice. Lysates were prepared by centrifugation at 16220 rpm for 10 minutes at 4°C and collecting the supernatant. The supernatant after centrifugation were then incubated with anti-Orai3 antibody (Abcam) and anti-MARCH8 antibody (Sigma) for reverse Co-IP at 4°C on an end-to-end rotor for overnight. Next day, the incubated lysates were precipitated by addition of activated protein A agarose beads for overnight at 4°C. The protein A agarose beads used for immunoprecipitation were first equilibrated with glycerol IP buffer and blocking was done using 0.1 µg/µl bovine serum albumin. Precipitations were also carried by IgG (Cell Signalling Technologies) raised in same host as the isotype control. The antigen-antibody complex obtained were processed in NP 40 lysis buffer for immunoblotting to check MARCH8 and Orai3 protein levels.

### Colocalization Immunofluorescence Assay

PANC-1 cells cultured on glass slides were transfected with eYFP-Orai3 using jetPRIME transfection reagent (114-01, Polyplus) as per the manufacturer’s protocol. After 48 h, the cells were washed with PBS and fixed with 4% paraformaldehyde in PBS for 15 mins. After three washes with PBS, the cells were permeabilized with PBS containing 0.3% Triton-X100 for 15 mins and blocked with PBS containing 2% bovine serum albumin and 0.1% Triton-X100 for 1 hr. The cells were then incubated with anti-MARCH8 antibody (1:200) for overnight. After incubation, the cells were washed thrice with PBS containing 0.1% Tween20 and incubated with Alexa Flour 568 goat anti rabbit-conjugated secondary antibody (1:500) at 25°C for 2 h. Finally, the cells were washed three times with PBS containing 0.1% Tween20 mounted using SlowFade Gold antifade reagent with 4′,6-diamidono-2-phenylindole (DAPI) (Thermo Fisher Scientific, Catalog #S36938) and visualized using a Carl Zeiss LSM 880 laser scanning confocal microscope at 63X (with oil) magnification.

### Methylated DNA immunoprecipitation (MeDIP)

The genomic DNA of MiaPaCa-2, PANC-1 and CFPAC-1 cells was extracted using DNeasy Blood & Tissue Kit (Catalog #69504) according to the manufacturer’s instructions. Genomic DNA was sonicated using Bioruptor Pico with 30s ON and 30s OFF for 10 cycles at 4°C to get DNA fragments between size 300–500 bp. Sonicated DNA was denatured by incubating at 95°C for 10 minutes followed by incubation on ice for 5 mins. 3 µg of sonicated DNA was incubated with 1 µg anti-5-Methylcytosine (Cell Signalling Technologies) antibody along with normal rabbit IgG at 4°C overnight. 30 µl magnetic beads were added and incubated overnight at 4°C. The beads were washed thrice with glycerol IP buffer at 4°C for 5 minutes and eluted in 150 µl elution buffer (1M Tris-HCl pH 8.0, 0.5M EDTA, 10% SDS) with proteinase K (Thermo Fisher Scientific, Catalog #EO0491) treatment for 2 h at 55°C on a rotating platform. Purification of DNA using phenol-chloroform extraction was performed and used for MeDIP-qPCR using primers specific to human MARCH8 core promoter. The primers used for MeDIP analysis are as follows:

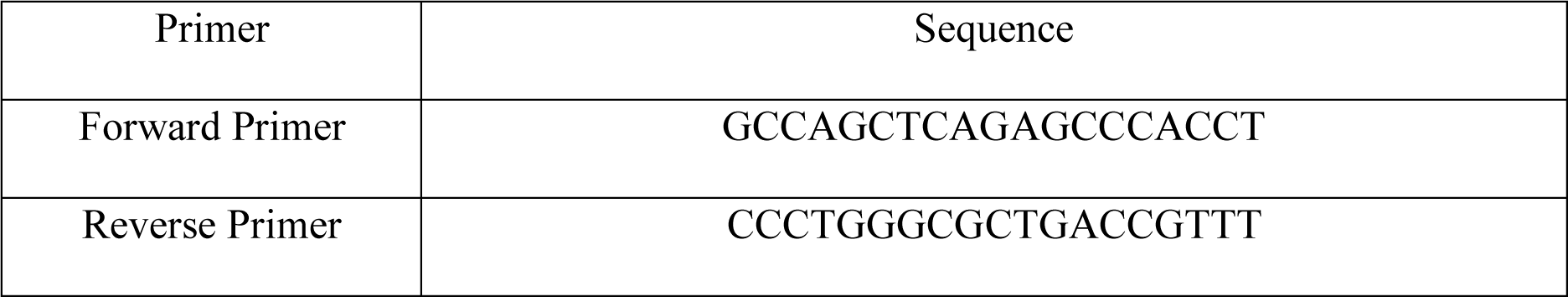

### Lentiviral Stable Cell Line Generation

For stable knockdown generation, human specific shNT and shMARCH8 cloned in the lentiviral pGIPZ vector (Dharmacon, Lafayette, CO, USA) were used. As reported earlier, the lentiviral constructs pCMV-VSVG, pCMV-dR8.2 and pGIPZ-shNT/shMARCH8 were co-transfected in a flask containing 95% confluent HEK-293T cells. Lipofectamine 2000 (Thermo Fisher, Waltham, MA, USA) was used as a transfection reagent to transfect HEK-293T cells. Viral particles containing medium were collected at 48 and 72 h after transfection and was concentrated using Amicon filters through centrifugation. These concentrated viral particles were used to transduce CFPAC-1 cells seeded at 50% confluency and knockdown was confirmed by performing Western blot analysis.

### Scratch wound healing assay

CFPAC-1 lentiviral stable cells were plated in a 24-well plate at a density of 3 x 10^5 cells per well. Once the cells reached full confluency, a scratch was made in the centre of each well using a P20 pipette tip. The cells were then rinsed with PBS, and fresh media was added to each well. Scratch wounds were monitored at various time points under a bright field microscope to observe cell migration. The migration rate of CFPAC-1 shNT and shMARCH8 cells was determined by measuring the distance travelled by the cells from the initial to the final time point. The wound closure was normalized to the 0-hour timepoints.

### Zebrafish Husbandry

The zebrafish used in this study were housed at the Regional Centre for Biotechnology (RCB). The zebrafish experiments were performed with the ethical approvals of the Institutional Animal Ethics Committee (IAEC), RCB. The reference number of the approval is RCB/IAEC/2022/143.

### Zebrafish Xenograft Microinjections

CFPAC-1 shNT and shMARCH8 knockdown cells were microinjected into the perivitelline space (PVS) of anesthetized 2 days post fertilization (dpf) zebrafish embryos and incubated at 34°C. 3 h post injection, the embryos were screened and visualized for metastasized events, the images were acquired in Nikon SMZ800N Stereo Microscope and injections were performed using Eppendorf FemtoJet 4i.

### Statistical Analysis

All the experiments were performed at least three times. Data are presented as mean ±SEM and one sample t-test was performed for determining the statistical significance. For experiments with more than two conditions, one-way ANOVA test was performed. p-value <0.05 was considered as significant and is presented as “*”; p-value < 0.01 is presented as “**”, p-value < 0.001 is presented as “***” and p-value < 0.0001 is presented as “****”.

## Acknowledgements

This work was supported by the Department of Biotechnology, India (project number BT/PR52477/MED/30/2540/2024). RKM also acknowledges funding support from RCB Institutional Core funding, Anusandhan National Research Foundation (ANRF) project number SERB-CRG/2023/004054 and DBT/Wellcome Trust India Alliance Fellowship (IA/I/19/2/504651). The authors thank members of the Motiani laboratory for discussions and critical reading of the manuscript. We thank Hideaki Fujita (Nagasaki International University, Japan) for sharing MARCH8-eGFP-N1 WT and MARCH8-eGFP-N1 CS plasmids. We also thank Mohamed Trebak (University of Pittsburg, US) for sharing Orai3-eYFP plasmid. The technical assistance of Mr. Unni Narayanan is highly appreciated.

## Authors Contributions

Sharon Raju: Methodology, Investigation, Visualization, Formal analysis, Writing-Original draft preparation. Akshay Sharma: Methodology, Investigation, Formal analysis, Visualization. Gyan Ranjan: Investigation, Visualization, Formal analysis. Rajender K Motiani: Conceptualization, Supervision, Writing-Original draft preparation, Reviewing and Editing, Project administration, Funding acquisition.

## Competing interests

Authors declare that they have no competing interests.

## Supplementary Figures

**Supplementary Figure 1:**
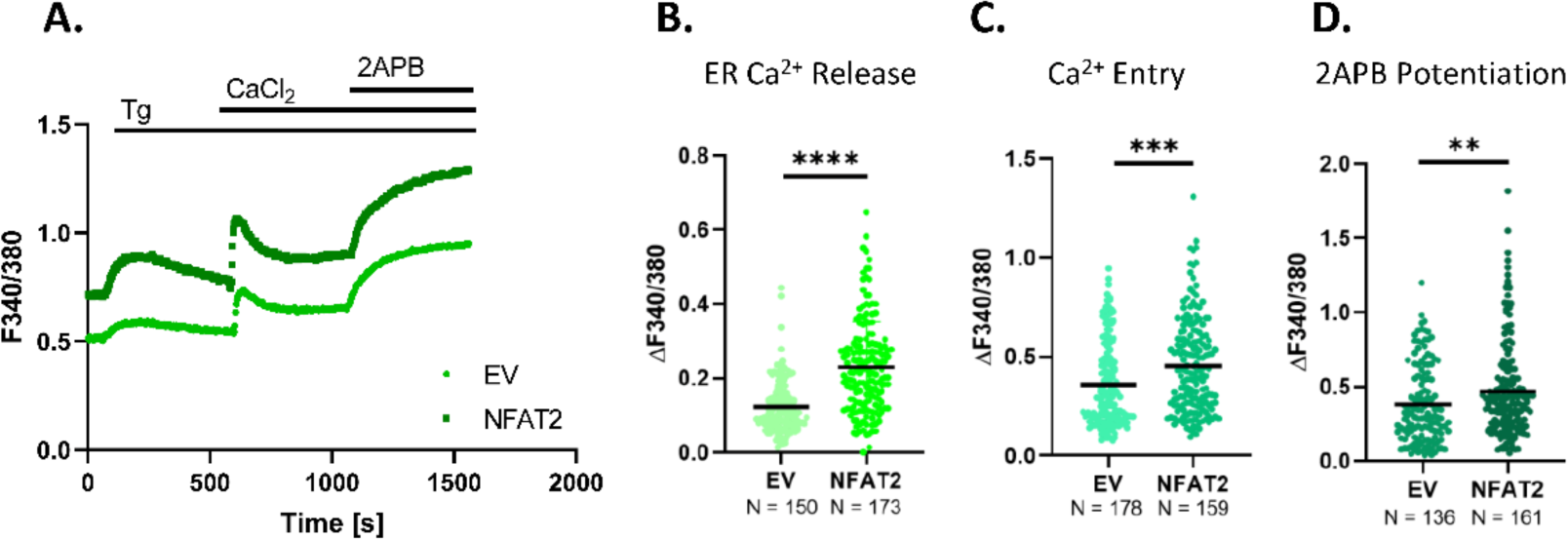
**A).** Representative Ca^2+^ imaging traces of empty vector control and NFAT2 overexpression in MiaPaCa-2. **B).** Quantitation of ER Ca^2+^release of NFAT2 overexpression in MiaPaCa-2 compared to empty vector control where “n” denotes the number of ROIs. **C).** Change in Ca^2+^entry upon overexpression of NFAT2 compared to empty vector control in MiaPaCa-2 where “n” denotes the number of ROIs. **D).** Orai3 potentiation of 2-APB in NFAT2 overexpressed and empty vector control MiaPaCa-2 where “n” denotes the number of ROIs. Data presented are mean ± S.E.M. For statistical analysis, unpaired student’s *t*-test was performed for panel B, C and D using GraphPad Prism software. Here, ** *p* < 0.01; *** *p* < 0.001 and **** *p* < 0.0001.

**Supplementary Figure 2:**
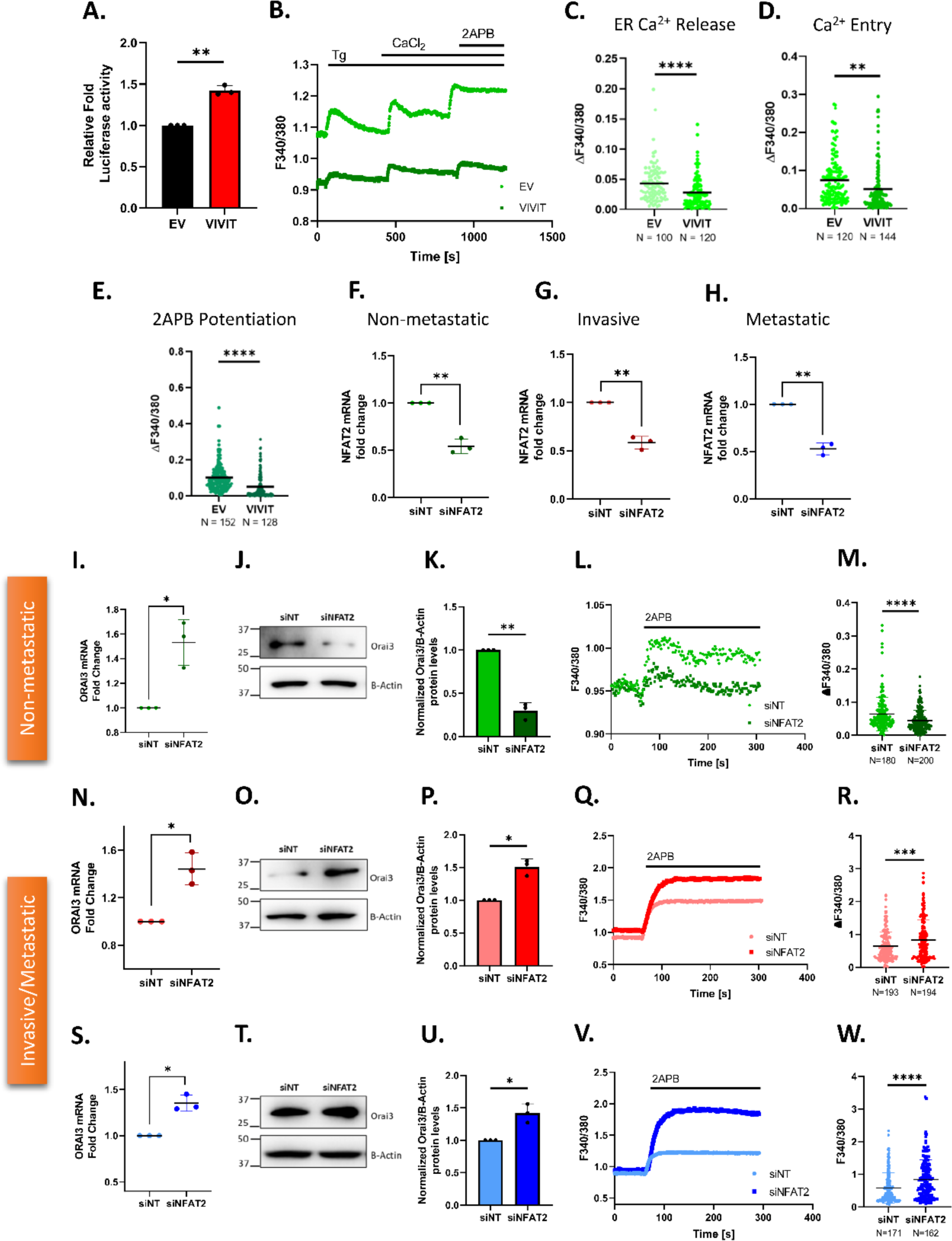
**A).** Normalized Luciferase activity of Orai3 promoter in PANC-1 cells upon VIVIT transfection for 48 hrs. (N=3). **B).** Representative Ca^2+^ imaging trace of control vector pEGFP-N1 plasmid and VIVIT transfection in MiaPaCa-2. **C).** Quantitation of ER Ca^2+^release of VIVIT Transfection in MiaPaCa-2 compared to empty vector control where “n” denotes the number of ROIs. **D).** Change in Ca^2+^entry upon VIVIT Transfection in MiaPaCa-2 where “n” denotes the number of ROIs. **E).** Orai3 potentiation of 2-APB in VIVIT Transfected and empty vector control MiaPaCa-2 where “n” denotes the number of ROIs. **F).** qRT-PCR analysis showing NFAT2 knockdown validation in MiaPaCa-2 (N=3). **G).** qRT-PCR analysis showing NFAT2 knockdown validation in PANC-1 (N=3). **H).** qRT-PCR analysis showing NFAT2 knockdown validation in CFPAC-1 (N=3). **I).** qRT-PCR analysis showing increase in Orai3 mRNA levels upon NFAT2 knockdown in MiaPaCa-2 compared to control (N=3). **J).** Representative western blots for decrease in Orai3 protein levels due to NFAT2 knockdown in MiaPaCa-2 cells compared to control. **K).** Densitometric quantitation of Orai3 protein levels in NFAT2 knockdown MiaPaCa-2 compared to control (N=3). **L).** Representative Ca^2+^ imaging trace of control siNT and siNFAT2 transfected MiaPaCa-2. **M).** Orai3 potentiation of 2-APB in siNT and siNFAT2 transfected MiaPaCa-2 where “n” denotes the number of ROIs. **N).** qRT-PCR analysis showing increase in Orai3 mRNA expression upon NFAT2 knockdown in PANC-1 compared to control (N=3). **O).** Representative western blots for increase in Orai3 protein levels due to NFAT2 knockdown in PANC-1 compared to control. **P).** Western Blot densitometry of Orai3 protein levels in NFAT2 knockdown PANC-1 cells compared to control (N=3). **Q).** Representative Ca^2+^ imaging trace of control siNT and siNFAT2 transfected PANC-1 cells. **R).** Potentiation of Orai3 by 2-APB in control siNT and siNFAT2 transfected PANC-1 where “n” denotes the number of ROIs. **S).** qRT-PCR analysis showing increase in Orai3 mRNA expression upon NFAT2 knockdown in CFPAC-1 compared to control (N=3). **T).** Representative western blots for decrease in Orai3 protein levels due to NFAT2 knockdown in CFPAC-1 compared to control. **U).** Western Blot densitometry of Orai3 protein in NFAT2 knockdown CFPAC-1 cells compared to control (N=3). **V).** Representative Ca^2+^ imaging trace of control siNT and siNFAT2 transfected CFPAC-1 cells. **W).** Potentiation of Orai3 by 2-APB in siNT and siNFAT2 transfected CFPAC-1 where “n” denotes the number of ROIs. Data presented are mean ± S.E.M. For statistical analysis, unpaired student’s *t*-test was performed for panel C, D, E, M, R and W while one sample *t*-test was performed for panels F, G, H, I, K, N, P, S and U using GraphPad Prism software. Here, * *p* <0.05; ** *p* < 0.01; *** *p* < 0.001 and **** *p* < 0.0001.

**Supplementary Figure 3:**
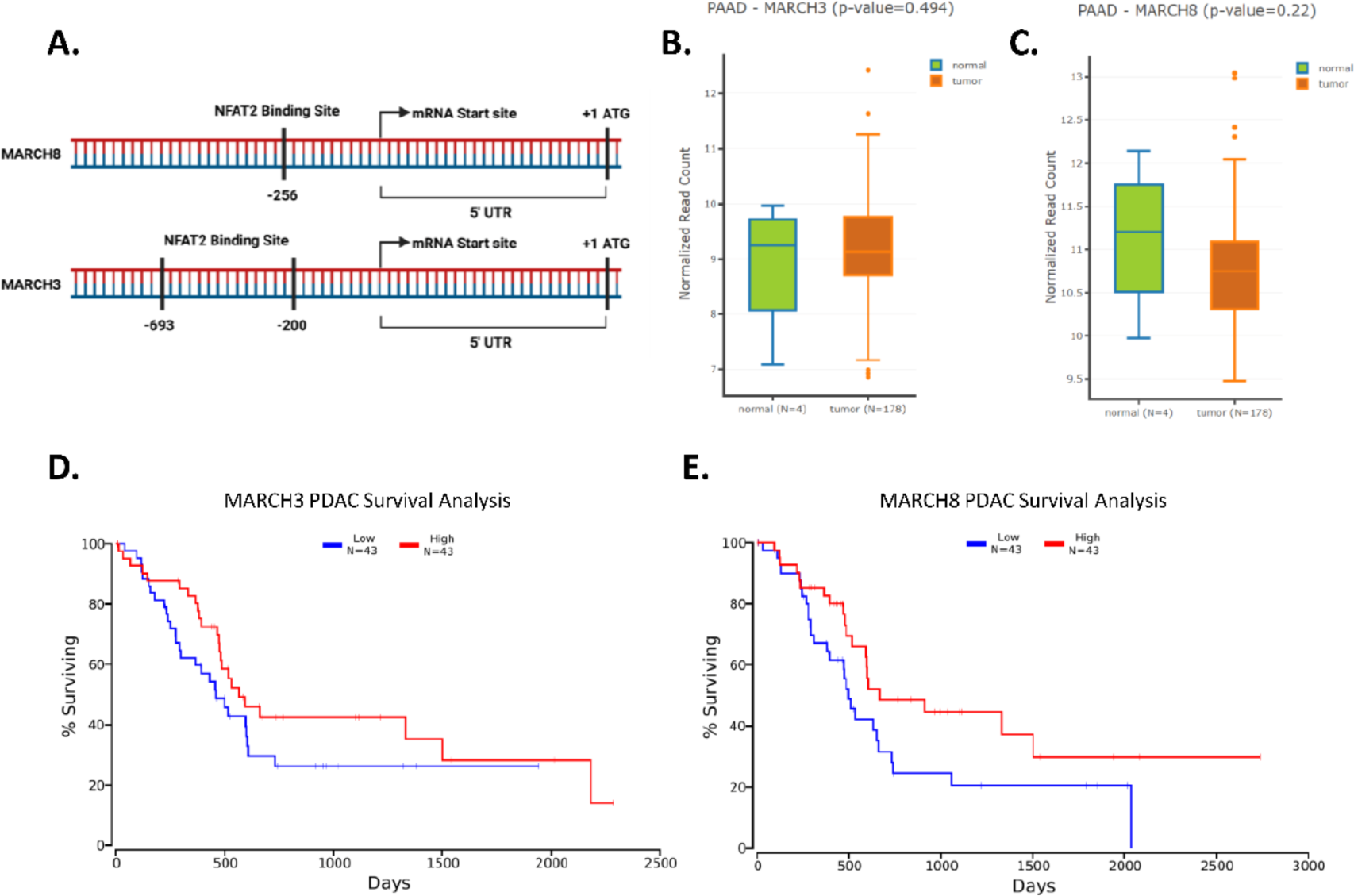
**A).** Identification of putative NFAT2 binding sites on the Human MARCH3 and MARCH8 promoter using the EPD-Search Motif Tool at p-val cut-off of 0.01. **B).** MARCH3 expression levels in normal pancreatic tissues and pancreatic adenocarcinoma (PAAD) tissues analysed by the DNMIVD database. **C).** MARCH8 expression levels in normal pancreatic tissues and pancreatic adenocarcinoma (PAAD) tissues analysed by the DNMIVD database. **D).** Survival Analysis in pancreatic cancer patients wherein blue trace corresponds to low MARCH3 expression (n=43), and red trace corresponds to high MARCH3 expression (n=43). **E).** Survival Analysis in pancreatic cancer patients wherein blue trace corresponds to low MARCH8 expression (n=43), and red trace corresponds to high MARCH8 expression (n=43).

**Supplementary Figure 4:**
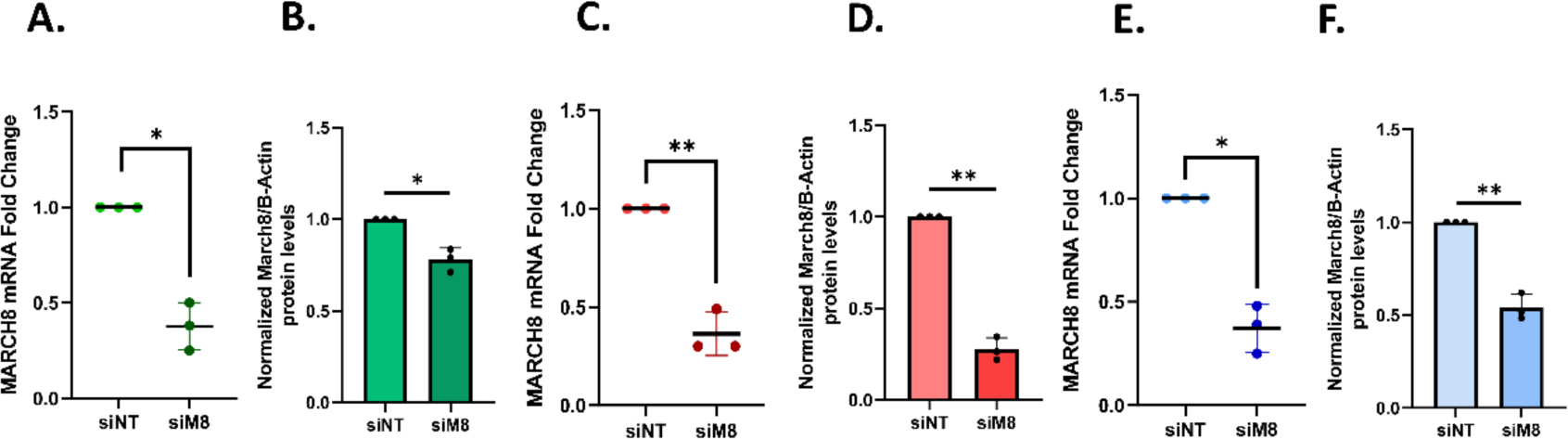
**A).** qRT-PCR analysis showing decrease in MARCH8 mRNA levels upon siMARCH8 knockdown in MiaPaCa-2 compared to control siNT (N=3). **B).** Densitometric quantitation of MARCH8 protein levels in siMARCH8 knockdown MiaPaCa-2 cells compared to siNT control (N=3). **C).** qRT-PCR analysis showing decrease in MARCH8 mRNA levels upon siMARCH8 knockdown in PANC-1 compared to control siNT (N=3). **D).** Densitometric quantitation of MARCH8 protein levels in siMARCH8 knockdown PANC-1 cells compared to siNT control (N=3). **E).** qRT-PCR analysis showing decrease in MARCH8 mRNA levels upon siMARCH8 knockdown in CFPAC-1 compared to control siNT (N=3). **F).** Densitometric quantitation of MARCH8 protein levels in siMARCH8 knockdown CFPAC-1 cells compared to siNT control (N=3). Data presented are mean ± S.E.M. For statistical analysis, one sample *t*-test was performed for panels A, B, C, D, E and F using GraphPad Prism software. Here, * *p* <0.05 and ** *p* < 0.01.

## Notes

### Competing Interest Statement

The authors have declared no competing interest.

### Summary of Updates

In this revised version, we have further validated our data with additional experiments. We have performed NFAT2 loss of function studies with siRNA mediated NFAT2 silencing as well. We have included this new data in the form of Supplementary Figure2. Also, we have updated patients' survival analysis data presented in the Figure9A and Supplementary Figure3D-E of this revised version. Finally, we have modified the title of the manuscript for better clarity.

## References

1. Anaya, J. (2016). OncoLnc: linking TCGA survival data to mRNAs, miRNAs, and lncRNAs. PeerJ Computer Science, 2, e67. 10.7717/peerj-cs.67

2. Arora, S., Tanwar, J., Sharma, N., Saurav, S., & Motiani, R. K. (2021). Orai3 regulates pancreatic cancer metastasis by encoding a functional store operated calcium entry channel. Cancers, 13(23). 10.3390/cancers13235937

3. Chalmers, S. B., & Monteith, G. R. (2018). ORAI channels and cancer. Cell Calcium, 74, 160–167. 10.1016/j.ceca.2018.07.011

4. Chen, C., Wang, Y., Zhao, Q., Li, G., Wang, Y., Xu, L., Huang, H., Song, G., Li, W., & He, X. (2023). E3 Ubiquitin Ligase MARCH8 Promotes Pancreatic Cancer Growth and Metastasis by Activating STAT3 via Degradation of PTPN4. Pancreas. 10.1097/MPA.0000000000002244

5. Chen, W., Patel, D., Jia, Y., Yu, Z., Liu, X., Shi, H., & Liu, H. (2021). MARCH8 Suppresses Tumor Metastasis and Mediates Degradation of STAT3 and CD44 in Breast Cancer Cells. Cancers, 13(11), 2550. 10.3390/cancers13112550

6. Ding, W., Chen, J., Feng, G., Chen, G., Wu, J., Guo, Y., Ni, X., & Shi, T. (2020). DNMIVD: DNA methylation interactive visualization database. Nucleic Acids Research, 48(D1), D856–D862. 10.1093/nar/gkz830

7. Fan, J., Tian, L., Li, M., Huang, S.-H., Zhang, J., & Zhao, B. (2017). MARCH8 is associated with poor prognosis in non-small cell lung cancers patients. Oncotarget, 8(64), 108238–108248. 10.18632/oncotarget.22602

8. Fitzmaurice, C., Abate, D., Abbasi, N., Abbastabar, H., Abd-Allah, F., Abdel-Rahman, O., Abdelalim, A., Abdoli, A., Abdollahpour, I., Abdulle, A. S. M., Abebe, N. D., Abraha, H. N., Abu-Raddad, L. J., Abualhasan, A., Adedeji, I. A., Advani, S. M., Afarideh, M., Afshari, M., Aghaali, M., … Murray, C. J. L. (2019). Global, regional, and national cancer incidence, mortality, years of life lost, years lived with disability, and disability-Adjusted life-years for 29 cancer groups, 1990 to 2017: A systematic analysis for the global burden of disease study. JAMA Oncology, 5(12), 1749–1768. 10.1001/jamaoncol.2019.2996

9. Fujita, H., Iwabu, Y., Tokunaga, K., & Tanaka, Y. (2013). Membrane-associated RING-CH (MARCH) 8 mediates the ubiquitination and lysosomal degradation of the transferrin receptor. Journal of Cell Science. 10.1242/jcs.119909

10. Gibney, E. R., & Nolan, C. M. (2010). Epigenetics and gene expression. Heredity, 105(1), 4–13. 10.1038/hdy.2010.54

11. Jones, P. A., & Taylor, S. M. (1980). Cellular differentiation, cytidine analogs and DNA methylation. Cell, 20(1), 85–93. 10.1016/0092-8674(80)90237-8

12. José Aramburu, Michael B. Yaffe, Cristina Lopez-Rodriguez, Lewis C. Cantley, Patrick G. Hogan, & Anjana Rao. (1999). Affinity driven peptide selection of an NFAT inhibitor more selective than cyclosporin A. Science.

13. Karolchik, D., Hinrichs, A. S., & Kent, W. J. (2009). The UCSC Genome Browser. *Current Protocols in Bioinformatics*, Chapter 1, Unit1.4. 10.1002/0471250953.bi0104s28

14. Kim, M.-H., Seo, J. B., Burnett, L. A., Hille, B., & Koh, D.-S. (2013). Characterization of store-operated Ca2+ channels in pancreatic duct epithelia. Cell Calcium, 54(4), 266–275. 10.1016/j.ceca.2013.07.002

15. Kreft, Ł., Soete, A., Hulpiau, P., Botzki, A., Saeys, Y., & De Bleser, P. (2017). ConTra v3: a tool to identify transcription factor binding sites across species, update 2017. Nucleic Acids Research, 45(W1), W490–W494. 10.1093/nar/gkx376

16. Lakshminarasimhan, R., & Liang, G. (2016). The Role of DNA Methylation in Cancer. Advances in Experimental Medicine and Biology, 945, 151–172. 10.1007/978-3-319-43624-1_7

17. Lee, D. H., & Goldberg, A. L. (1998). Proteasome inhibitors: valuable new tools for cell biologists. Trends in Cell Biology, 8(10), 397–403. 10.1016/S0962-8924(98)01346-4

18. Li, L.-C., & Dahiya, R. (2002). MethPrimer: designing primers for methylation PCRs. Bioinformatics, 18(11), 1427–1431. 10.1093/bioinformatics/18.11.1427

19. Lin, H., Li, S., & Shu, H.-B. (2019a). The Membrane-Associated MARCH E3 Ligase Family: Emerging Roles in Immune Regulation. Frontiers in Immunology, 10. 10.3389/fimmu.2019.01751

20. Lin, H., Li, S., & Shu, H.-B. (2019b). The Membrane-Associated MARCH E3 Ligase Family: Emerging Roles in Immune Regulation. Frontiers in Immunology, 10, 1751. 10.3389/fimmu.2019.01751

21. Lopez, J. J., Jardin, I., Albarrán, L., Sanchez-Collado, J., Cantonero, C., Salido, G. M., Smani, T., & Rosado, J. A. (2020). Molecular Basis and Regulation of Store-Operated Calcium Entry (pp. 445–469). 10.1007/978-3-030-12457-1_17

22. Martinez-Lopez, M., Póvoa, V., & Fior, R. (2021). Generation of Zebrafish Larval Xenografts and Tumor Behavior Analysis. Journal of Visualized Experiments, 172. 10.3791/62373

23. Motiani, R. K., Abdullaev, I. F., & Trebak, M. (2010). A Novel Native Store-operated Calcium Channel Encoded by Orai3. Journal of Biological Chemistry, 285(25), 19173–19183. 10.1074/jbc.M110.102582

24. Motiani, R. K., Zhang, X., Harmon, K. E., Keller, R. S., Matrougui, K., Bennett, J. A., & Trebak, M. (2013). Orai3 is an estrogen receptor α-regulated Ca2+ channel that promotes tumorigenesis. The FASEB Journal, 27(1), 63–75. 10.1096/fj.12-213801

25. Müller, M. R., & Rao, A. (2010a). NFAT, immunity and cancer: a transcription factor comes of age. Nature Reviews Immunology, 10(9), 645–656. 10.1038/nri2818

26. Müller, M. R., & Rao, A. (2010b). NFAT, immunity and cancer: a transcription factor comes of age. Nature Reviews Immunology, 10(9), 645–656. 10.1038/nri2818

27. Nishiyama, A., & Nakanishi, M. (2021). Navigating the DNA methylation landscape of cancer. Trends in Genetics, 37(11), 1012–1027. 10.1016/j.tig.2021.05.002

28. Perier, R. C. (2000). The Eukaryotic Promoter Database (EPD). Nucleic Acids Research, 28(1), 302–303. 10.1093/nar/28.1.302

29. Prakriya, M., Feske, S., Gwack, Y., Srikanth, S., Rao, A., & Hogan, P. G. (2006). Orai1 is an essential pore subunit of the CRAC channel. Nature, 443(7108), 230–233. 10.1038/nature05122

30. Qian, G., Guo, J., Vallega, K. A., Hu, C., Chen, Z., Deng, Y., Wang, Q., Fan, S., Ramalingam, S. S., Owonikoko, T. K., Wei, W., & Sun, S.-Y. (2021). Membrane-Associated RING-CH 8 Functions as a Novel PD-L1 E3 Ligase to Mediate PD-L1 Degradation Induced by EGFR Inhibitors. Molecular Cancer Research, 19(10), 1622– 1634. 10.1158/1541-7786.MCR-21-0147

31. Ross, A. B., Langer, J. D., & Jovanovic, M. (2021). Proteome Turnover in the Spotlight: Approaches, Applications, and Perspectives. Molecular & Cellular Proteomics, 20, 100016. 10.1074/mcp.R120.002190

32. Singh, S., Saraya, A., Das, P., & Sharma, R. (2017). Increased expression of MARCH8, an E3 ubiquitin ligase, is associated with growth of esophageal tumor. Cancer Cell International, 17(1), 116. 10.1186/s12935-017-0490-y

33. Tang, Z., Li, C., Kang, B., Gao, G., Li, C., & Zhang, Z. (2017). GEPIA: a web server for cancer and normal gene expression profiling and interactive analyses. Nucleic Acids Research, 45(W1), W98–W102. 10.1093/nar/gkx247

34. Tanwar, J., Ahuja, K., Sharma, A., Sehgal, P., Ranjan, G., Sultan, F., Agrawal, A., D’Angelo, D., Priya, A., Yenamandra, V. K., Singh, A., Raffaello, A., Madesh, M., Rizzuto, R., Sivasubbu, S., & Motiani, R. K. (2024). Mitochondrial calcium uptake orchestrates vertebrate pigmentation via transcriptional regulation of keratin filaments. PLOS Biology, 22(11), e3002895. 10.1371/journal.pbio.3002895

35. Tanwar, J., Arora, S., & Motiani, R. K. (2020). Orai3: Oncochannel with therapeutic potential. In Cell Calcium (Vol. 90). Elsevier Ltd. 10.1016/j.ceca.2020.102247

36. Tanwar, J., Sharma, A., Saurav, S., Shyamveer, Jatana, N., & Motiani, R. K. (2022). MITF is a novel transcriptional regulator of the calcium sensor STIM1: Significance in physiological melanogenesis. Journal of Biological Chemistry, 298(12), 102681. 10.1016/j.jbc.2022.102681

37. Tapper, H., & Sundler, R. (1995). Bafilomycin A1 inhibits lysosomal, phagosomal, and plasma membrane H+ -ATPase and induces lysosomal enzyme secretion in macrophages. Journal of Cellular Physiology, 163(1), 137–144. 10.1002/jcp.1041630116

38. Vaeth, M., & Feske, S. (2018). NFAT control of immune function: New Frontiers for an Abiding Trooper. F1000Research, 7, 260. 10.12688/f1000research.13426.1

39. Vashisht, A., Tanwar, J., & Motiani, R. K. (2018). Regulation of proto-oncogene Orai3 by miR18a/b and miR34a. Cell Calcium, 75, 101–111. 10.1016/j.ceca.2018.08.006

40. Vashisht, A., Trebak, M., & Motiani, R. K. (2015). STIM and Orai proteins as novel targets for cancer therapy. A Review in the Theme: Cell and Molecular Processes in Cancer Metastasis. American Journal of Physiology-Cell Physiology, 309(7), C457– C469. 10.1152/ajpcell.00064.2015

41. Wang, X., Li, Y., He, M., Kong, X., Jiang, P., Liu, X., Diao, L., Zhang, X., Li, H., Ling, X., Xia, S., Liu, Z., Liu, Y., Cui, C.-P., Wang, Y., Tang, L., Zhang, L., He, F., & Li, D. (2022). UbiBrowser 2.0: a comprehensive resource for proteome-wide known and predicted ubiquitin ligase/deubiquitinase–substrate interactions in eukaryotic species. Nucleic Acids Research, 50(D1), D719–D728. 10.1093/nar/gkab962

42. Wang, Z., Wang, M.-M., Geng, Y., Ye, C.-Y., & Zang, Y.-S. (2022). Membrane-associated RING-CH protein (MARCH8) is a novel glycolysis repressor targeted by miR-32 in colorectal cancer. Journal of Translational Medicine, 20(1), 402. 10.1186/s12967-022-03608-z

43. White, R. M., & Patton, E. E. (2023). Adult zebrafish as advanced models of human disease. Disease Models & Mechanisms, 16(8). 10.1242/dmm.050351

44. Xu, Y., Zhang, D., Ji, J., & Zhang, L. (2023). Ubiquitin ligase MARCH8 promotes the malignant progression of hepatocellular carcinoma through PTEN ubiquitination and degradation. Molecular Carcinogenesis, 62(7), 1062–1072. 10.1002/mc.23546

45. Ying, L., Li, K., Chen, C., Wang, Y., Zhao, Q., Wang, Y., Xu, L., Huang, H., Song, G., Li, W., & He, X. (2024). OIP5-AS1 enhances the malignant characteristics and resistance to chemotherapy of pancreatic cancer cells by targeting miR-30d-5p/MARCH8. Heliyon, 10(13), e33835. 10.1016/j.heliyon.2024.e33835

46. Zambelli, F., Pesole, G., & Pavesi, G. (2009). Pscan: finding over-represented transcription factor binding site motifs in sequences from co-regulated or co-expressed genes. Nucleic Acids Research, 37(suppl_2), W247–W252. 10.1093/nar/gkp464

47. Zhao, L., Zhao, J., Zhong, K., Tong, A., & Jia, D. (2022). Targeted protein degradation: mechanisms, strategies and application. Signal Transduction and Targeted Therapy, 7(1), 113. 10.1038/s41392-022-00966-4

